# Evolution of HMA-integrated tandem kinases accompanied by expansion of target pathogens

**DOI:** 10.64898/2025.12.15.692859

**Authors:** Soichiro Asuke, Analiza G. Tagle, Gang-Su Hyon, Sayako Koizumi, Tsubasa Murakami, Akiko Horie, Daisuke Niwamoto, Emi Katayama, Mai Shibata, Yoshino Takahashi, M T. Islam, Yoshihiro Matsuoka, Nami Yamaji, Motoki Shimizu, Ryohei Terauchi, Hiroshi Hisano, Kazuhiro Sato, Yukio Tosa

## Abstract

Tandem kinase proteins (TKPs) are an emerging family of plant intracellular immune receptors that offer potential for developing novel disease resistance. We cloned *Rmo2* and *Rwt7*, genes for resistance to the blast fungus, from barley and wheat, respectively, and discovered that they are orthologs encoding TKPs with an integrated N-terminal heavy metal-associated (HMA) domain. *Rmo2* was collocated with *Rpg1*, a TKP gene for resistance to stem rust, while *Rwt7* was collocated with *WTK4*, a TKP gene for resistance to powdery mildew. Domain swapping suggested that the HMA domain determines target effector specificity of these genes. We propose a model illustrating an evolutionary process in which a TKP gene has differentiated into paralogs and orthologs that recognize various effectors through diversification of their HMA domains.

## Main Text

Plants defend themselves against microbes and pests through a two layered innate immune system: pattern-triggered immunity (PTI) and effector-triggered immunity (ETI) (*1*). ETI is initiated upon recognition of pathogen effectors (products of avirulence genes) by products of resistance genes. Most resistance genes cloned to date encode nucleotide-binding leucine-rich repeat (NLR) proteins (*2–4*). NLRs recognize corresponding effectors either through direct binding or by monitoring modifications of effector targets or decoy proteins that mimic the targets (*4–7*). Some resistance genes comprise paired NLRs, where one encodes a sensor NLR protein and the other encodes a helper NLR protein (*7*). Analyses of paired NLR systems led to the establishment of the integrated decoy model, in which decoy domains are integrated into NLR receptors to trap effectors directly (*8–10*). For example, the rice NLR proteins RGA5 and Pik-1, encoded by the rice blast resistance genes *Pia* (*11*) and *Pik* (*12, 13*), respectively, contain heavy metal-associated (HMA) domains that trap rice blast effectors and trigger defense responses (*14, 15*).

Recently, tandem kinase proteins (TKPs), which possess two kinase and/or pseudokinase domains, have emerged as another important family of ETI regulators (*2, 16*). The first isolated resistance gene encoding TKP was *Rpg1*, which confers resistance in barley to the stem rust fungus (*17, 18*). *Rpg1* encodes a protein with a pseudokinase domain (PKD) – kinase domain (KD) architecture. To date, ∼10 resistance genes encoding TKPs have been isolated from Triticeae species (*19, 20*). Recent studies have suggested that products of *Sr62* and *Rwt4*, TKP genes with a KD-PKD architecture, trap their corresponding effectors via their N-terminal kinase and partially duplicated kinase fragment, respectively (*21–23*). These studies also demonstrated that the same helper NLR is involved in the resistance mediated by *Sr62* and *Rwt4* (*21, 22*).

*Pyricularia oryzae* (syn. *Magnaporthe oryzae*), a causal agent of blast diseases in gramineous plants, consists of several host genus-specific pathotypes such as MoO (*<u>M</u>. <u>o</u>ryzae* pathotype *<u>O</u>ryza*), MoS, MoE, MoL, MoA, and MoT, which are pathogenic on *Oryza*, *Setaria*, *Eleusine*, *Lolium*, *Avena*, and *Triticum*, respectively (*24–26*). MoE is further divided into two subgroups, EC-I and EC-II (*27*). MoT (the wheat blast fungus) first emerged in Brazil in 1985 (*28*) and is now causing a pandemic disease (*29*). MoT is suggested to have evolved from MoL or its relatives through the loss of function of the avirulence gene *PWT3*, which corresponds to the resistance gene *Rwt3* (*30*). To further investigate the history of pathotype differentiation, we performed genetic analyses with MoE and MoT, and found that at least five genes are involved in the avirulence of MoE (EC-II) on wheat (*31*). One of the five genes was an allele of *PWT3* (*31*). Subsequently, we identified the second avirulence gene, *PWT6* (*32*), along with its corresponding resistance gene *Rwt6* (*33*). Furthermore, we found that the third avirulence gene was a homolog of *PWT7* involved in the avirulence of an *Avena* isolate on wheat (*34*). However, its corresponding resistance gene, tentatively designated as *Rwt7*, has not been identified.

*P. oryzae* does not contain a pathotype specialized for barley. When barley cultivars are inoculated with the pathotypes mentioned above, they exhibit various reactions similar to those observed in race-cultivar specificity (*35*). Although each cultivar shows resistance to a different range of pathotypes/isolates, all of the resistance is attributed to a single locus designated as *RMO2* (*36*). These cultivars carry different *Rmo2* alleles that recognize different sets of avirulence genes. These results suggest that *RMO2* is a core locus governing barley resistance to *P. oryzae*.

In the present study we cloned *Rmo2* from barley and *Rwt7* from wheat, and found that they are orthologous genes encoding TKPs with a HMA domain. We also cloned the avirulence gene *PBY2* corresponding to *Rmo2.d*. Based on interactions between these resistance genes and the avirulence genes (*PWT7* and *PBY2*), we propose a model illustrating an evolutionary process of these TKP genes.

### Cloning of *RMO2*

To clone *RMO2*, we performed infection assays with representative barley cultivars (Fig. 1A). ‘Russia 81’(R81) is resistant to all of three isolates, GFSI1-7-2 (MoS), MZ5-1-6 (MoE), and Br48 (MoT). However, ‘Russia 74’(R74) is resistant to GFSI1-7-2 and MZ5-1-6 but susceptible to Br48, and ‘H.E.S.4’(H4) is resistant to GFSI1-7-2 alone. The reactions of R81, R74, and H4 are controlled by different alleles, *Rmo2.d*, *Rmo2.c*, and *Rmo2.a*, respectively, located at the same genetic locus *RMO2* (*36*). We crossed R81 with ‘Nigrate’(Ngt) carrying *rmo2* (Fig. 1A), inoculated resulting 93 F_2:3_ lines with GFSI1-7-2, and identified some flanking markers through rough mapping (Fig. 1B). For fine mapping, a total of ∼26,000 F_3_ individuals from segregating F_2:3_ lines were screened for recombinants between flanking markers. *RMO2* genotypes of the recombinants were determined by inoculation with GFSI1-7-2 in the next generation (F_4_). Consequently, the candidate region harboring *RMO2* was narrowed down to 62 kb flanked by AGT143 and AGT125 (Fig. 1C) on the Morex V3 reference genome. This region contained only one high confidence gene, *ABC1037* (*37*) encoding a TKP with a PKD-KD architecture preceded by an N-terminal HMA domain (Fig. 1D, E).

**Fig. 1.**
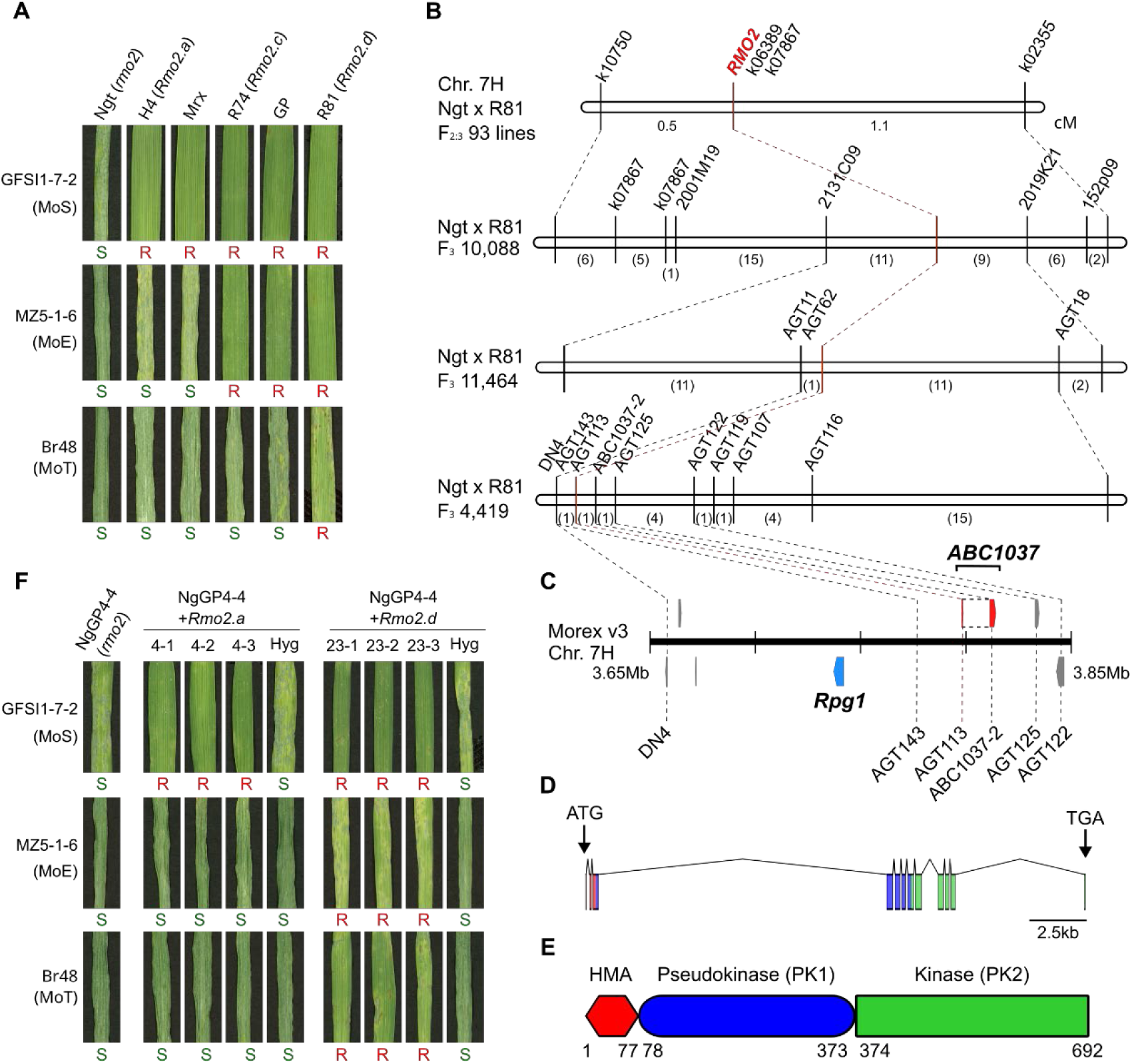
Cloning of *RMO2*, a core locus conditioning reactions of barley to *P. oryzae*. (**A**) Reactions of barley cultivars ‘Nigrate’ (Ngt), ‘H.E.S.4’ (H4), ‘Morex’ (Mrx), ‘Russia 74’ (R74), ‘Golden Promise’ (GP), and ‘Russia 81’ (R81) to *P. oryzae* isolates GFSI1-7-2, MZ5-1-6, and Br48. Genotypes of the cultivars and pathotypes of the isolates are shown in parentheses. R, resistant; S, susceptible. (**B**) Genetic maps around *RMO2* on chromosome 7H constructed using F_2:3_ lines and their descendants derived from an Ngt × R81 cross. Genotypes at the *RMO2* locus were determined by inoculation with GFSI1-7-2. Rough mapping with the F_2:3_ lines was followed by three rounds of fine mapping with F_3_ plants derived from heterozygous F_2_ lines. In each round, the segregating F_3_ populations were screened for recombinants between the far left and far right markers. Numbers of recombinants between internal markers are shown in the parentheses. (**C**) Physical map of the *RMO2* region based on the Morex V3 reference genome. The red and blue boxes indicate *ABC1037* (the *Rmo2* candidate gene) and *Rpg1* (a gene for resistance to *Puccinia graminis* f.sp. *tritici*), respectively. (**D**) Structure of the *ABC1037* gene. (**E**) Structure of ABC1037 in R81 (carrying *Rmo2.d*). (**F**) Recapitulation of the specificity of *Rmo2* alleles against three pathotypes of *P. oryzae*. NgGP4-4 (the recipient barley line) and T_2_ plants derived from its transformation with cDNAs of the *Rmo2.a* and *Rmo2.d* candidate genes (*ABC1037* in H4 and R81) were inoculated with GFSI1-7-2 (MoS), MZ5-1-6 (MoE), and Br48 (MoT). T_2_ lines carrying the *Rmo2.a* candidate (4–1, 4–2, 4–3) and the *Rmo2.d* candidate (23–1, 23–2, 23–3) are homozygous for the transgenes. Hyg represents a T_2_ line harboring no transgene.

*ABC1037* was polymorphic between R81 (*Rmo2.d*) and Ngt (*rmo2*) (fig. S1), although it was expressed in both (fig. S2). Furthermore, the *RMO2* phenotype perfectly co-segregated with a marker designed on its HMA domain (AGT113) but had one recombinant with a marker designed on its kinase domain (ABC1037-2) (Fig. 1B). This recombinant displayed intragenic recombination within *ABC1037* and lost the resistance to GFSI1-7-2. These results strongly suggested that *ABC1037* is *RMO2*. Interestingly, *Rpg1* was located 56 kb upstream of *ABC1037* on the Morex V3 reference genome (Fig. 1C).

To confirm the function of *ABC1037 in vivo*, we performed transformation of barley. Unfortunately, ‘Golden Promise’ (GP), the highly transformable barley cultivar widely used in barley genetic modification, was resistant to GFSI1-7-2 and MZ5-1-6 (Fig. 1A). Therefore, a transformable barley line carrying *rmo2* (NgGP4-4) was produced through screening descendants from Ngt x GP (fig. S3). *ABC1037* was cloned from transcripts of R81 (*Rmo2.d*) and H4 (*Rmo2.a*) (fig. S1) and introduced into NgGP4-4. T_2_ plants carrying the R81-derived *ABC1037* (homozygous) were resistant to all of the three isolates, while those carrying the H4-derived *ABC1037* were resistant to GFSI1-7-2 alone (Fig. 1F). This indicates that the transformation with *ABC1037* phenocopied the resistance pattern conferred by the *Rmo2* alleles (Fig. 1A). From these results, we conclude that *RMO2* is *ABC1037*.

### Cloning of *RWT7*

To identify *Rwt7*, we performed infection assays of representative wheat cultivars (Fig. 2A). ‘Norin 4’(N4), ‘Chinese Spring’(CS), ‘Hope’, and ‘Transfed’(Tfed) are susceptible to Br48 (MoT). Against Br48+PWT7 (a transfomant of Br48 carrying the *PWT7* transgene), however, N4, CS, and Hope are resistant but Tfed is susceptible, suggesting that the putative resistance gene (*Rwt7*) corresponding to *PWT7* is carried by N4, CS, and Hope but not by Tfed. Genetic analyses confirmed that the resistance of N4, CS, and Hope is controlled by the same, single gene (table S1). This gene was designated as *Rmg12* (with *Rwt7* listed as a synonym).

**Fig. 2.**
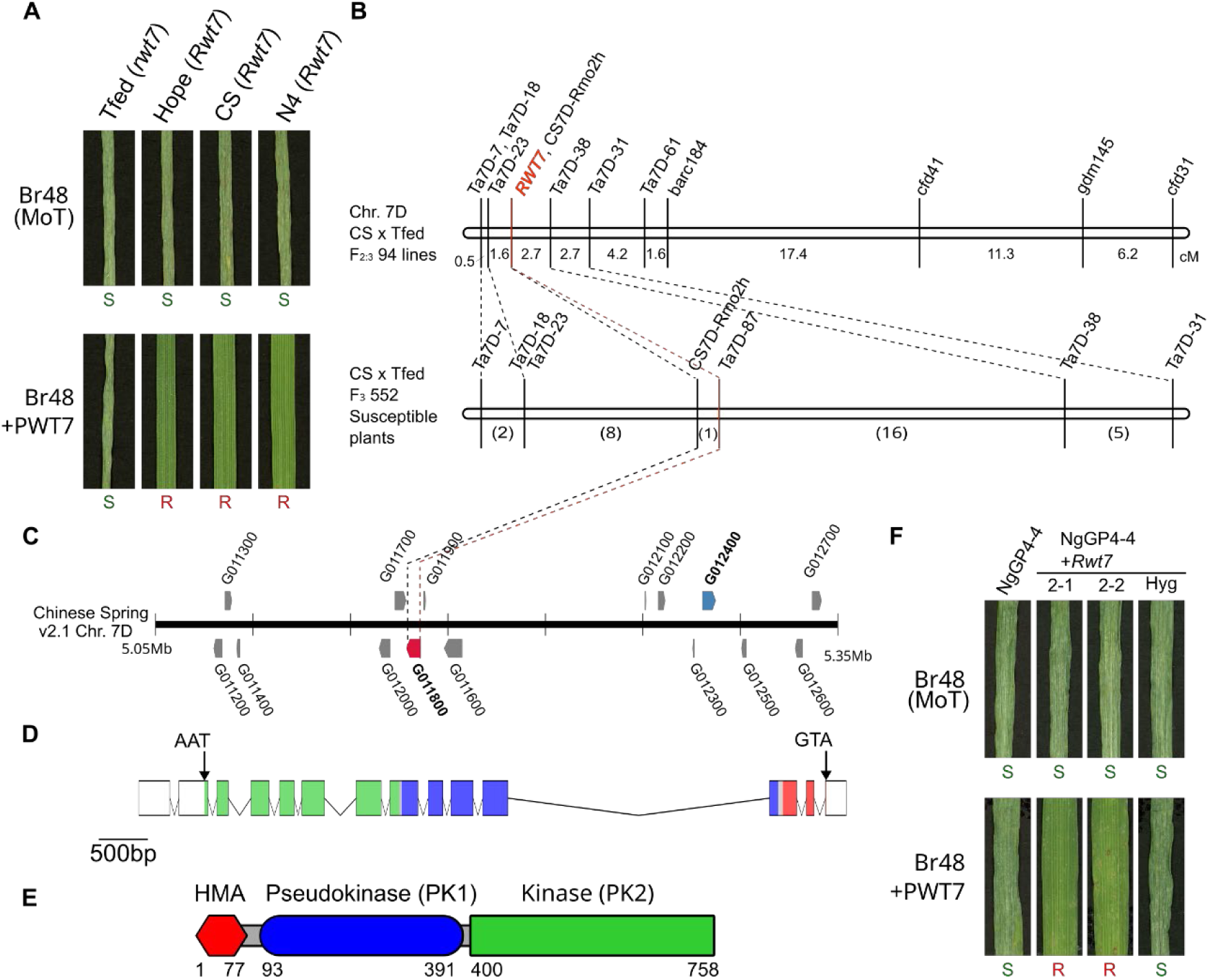
Cloning of *RWT7*, a fundamental locus conditioning the incompatibility between non-adapted pathotypes of *P. oryzae* and common wheat. (**A**) Reactions of common wheat cultivars ‘Transfed’ (Tfed), ‘Hope’, and ‘Chinese Spring’ (CS) to Br48 and Br48+PWT7 (a transformant of Br48 carrying *PWT7*). Genotypes of the cultivars and the pathotype of the isolate are shown in parentheses. R, resistant; S, susceptible. (**B**) Genetic maps around *RWT7* on chromosome 7D constructed using F_2:3_ lines derived from CS × Tfed. Rough mapping with theF_2:3_ lines was followed by fine mapping with F_3_ plants derived from heterozygous F_2_ lines. In the rough mapping, genotypes at the *RWT7* locus were determined by reactions of F_2:3_ lines to Br48+PWT7. In the fine mapping, the segregating F_3_ populations were first inoculated with Br48+PWT7, then resulting susceptible plants were screened for recombinants between the far left and far right markers. Numbers of recombinants between internal markers are shown in the parentheses. (**C**) Physical map of the *Rwt7* region based on the Chinese Spring v2.1 reference genome. The red and blue boxes indicate G011800 (the *Rwt7* candidate gene) and G012400 (an ortholog of *Rpg1*), respectively. (**D**) Structure of G011800. (**E**) Structure of the G011800 product. (**F**) Reactions of transformants carrying *Rwt7* to Br48 and Br48+PWT7. NgGP4-4 and its T_2_ transformants carrying cDNA of G011800 (the *Rwt7* candidate gene) were inoculated with Br48 and Br48+PWT7. 2-1 and 2-2 are T_2_ lines homozygous for the transgene. Hyg represents a T_2_ line harboring no transgene.

To clone *Rwt7*, we produced 94 F_2:3_ lines derived from CS x Tfed. Rough mapping with these F_2:3_ lines revealed that *Rwt7* is located on a region on chromosome 7D syntenic to barley *RMO2* (Fig. 2B). For fine mapping, a total of 3069 F_3_ individuals from the segregating F_2:3_ lines were inoculated with Br48+PWT7, and 552 susceptible individuals were subjected to screening for recombinants between flanking markers. The delimited candidate region contained TraesCS7D02G011800 (abbreviated as G011800 hereafter) (Fig. 2C) encoding a TKP with a PKD-KD architecture preceded by an HMA domain (Fig. 2D, E) on the Chinese Spring v2.1 reference genome. G011800 was expressed in CS but not in Tfed (fig. S4). To confirm its function *in vivo*, G011800 was cloned from transcripts of CS and introduced into NgGP4-4. T_2_ plants carrying the G011800 transgene (homozygous) were resistant to Br48+PWT7 but susceptible to Br48 (Fig. 2F), suggesting that G011800 specifically recognizes *PWT7* and therefore is *Rwt7*. Interestingly, another TKP gene (TraesCS7D02G012400, abbreviated as G012400, hereafter) with a PKD-KD architecture was located on 144 kb upstream of *Rwt7* (Fig. 2C).

### Cloning of *PBY2* and *PBY1*

R81 is resistant to both Br48 and MZ5-1-6 (Fig. 1A), but different avirulence genes underpin these interactions. *Rmo2.d* in R81 recognizes *PBY2* of Br48 and *PBY1* of MZ5-1-6 (*36*). To isolate *PBY2*, 200R45 and 200R69, F_1_ cultures virulent on R81, were backcrossed with Br48, resulting in a total of 86 BC_1_F_1_ cultures in which *PBY2* alone was segregating (Fig. 3A).

**Fig. 3.**
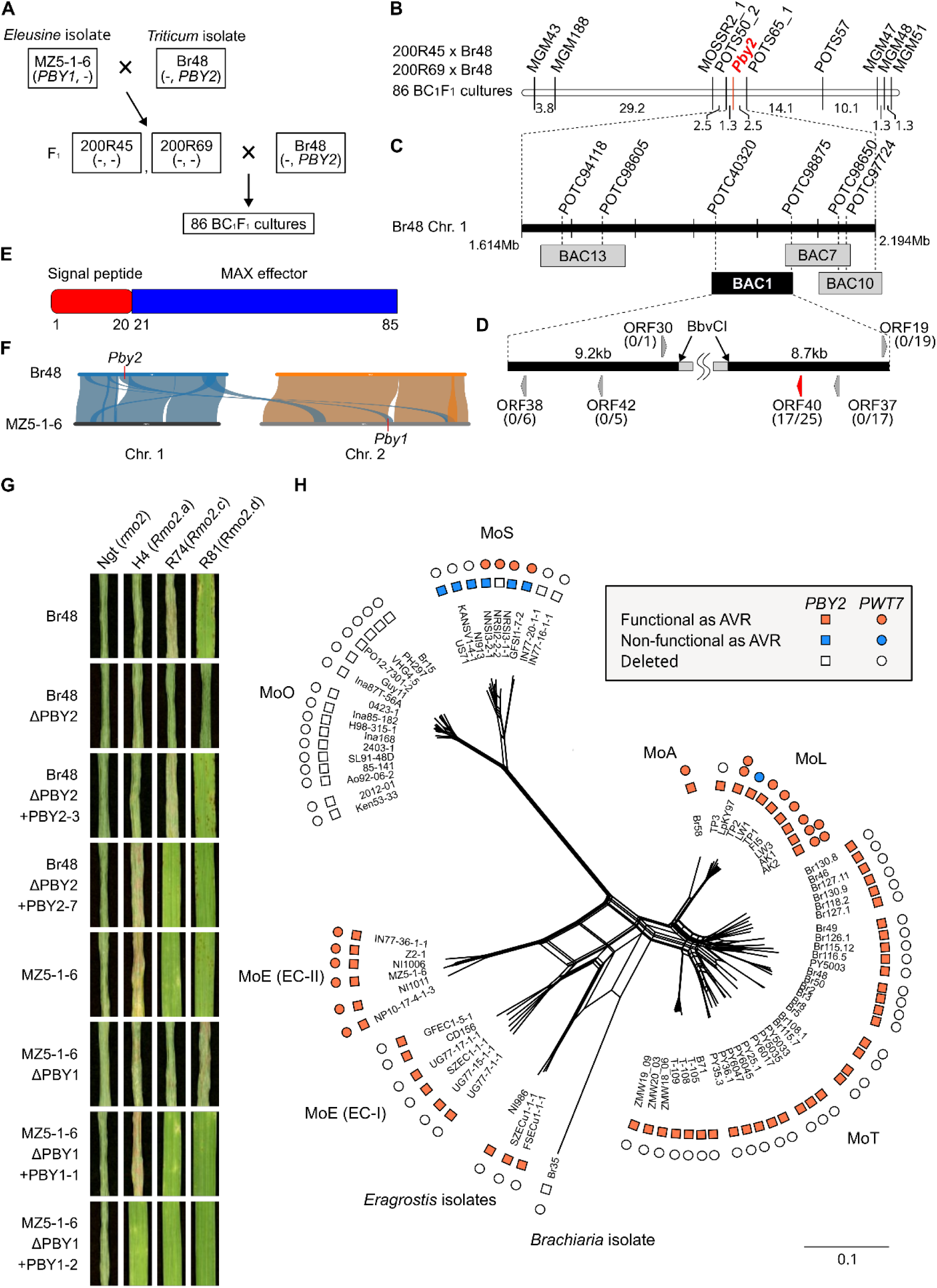
Cloning of *PBY2*, an avirulence gene conditioning the incompatibility between various pathotypes of *P. oryzae* and barley. (**A**) Pedigree of *P. oryzae* strains used for mapping of *PBY2*. (**B**) Genetic maps around *PBY2* on chromosome 1 of Br48 constructed using 86 BC_1_F_1_ cultures derived from two backcrosses, 200R45 × Br48 and 200R69 × Br48. Genotypes at the *Pby2* locus were determined by reactions of the BC_1_F_1_ cultures on R81. (**C**) Physical map around *Pby2* on chromosome 1 of Br48 along with BAC clones covering the candidate region. (**D**) Structure of BAC1 which conferred avirulence on R81 to 200R45 in a transformation assay. Arrowheads indicate putative effector genes predicted with SignalP4.1. They were cloned into pBluescript II SK(+) and introduced into 200R45. Shown in parentheses are numbers of transformants avirulent on R81/ numbers of transformants tested. (**E**) Structure of *PBY2* (ORF40). (**F**) Comparison of chromosomes 1 and 2 of MZ5-1-6 and Br48. *PBY2* represents ORF40 in Br48 (ORF40^Br^) while *PBY1* represents a gene identical to ORF4 in MZ6-1-6 (ORF4^MZ^) (**G**) Validation of the function of *PBY2* and *PBY1.* Eight-day-old primary leaves of Ngt, H4, R74, and R81 were inoculated with Br48 (MoT), Br48ΔPBY2 (Br48 with disrupted ORF40^Br^), Br48ΔPBY2+PBY2 (Br48ΔPBY2 carrying re-introduced ORF40^Br^), MZ5-1-6 (MoE), MZ5-1-6ΔPBY1 (MZ5-1-6 with disrupted ORF40^MZ^), and MZ5-1-6ΔPBY1+PBY1 (MZ5-1-6ΔPBY1 carrying re-introduced ORF40 ^MZ^). Genotypes of the cultivars are shown in parentheses. (**H**) Distribution of *PBY2* and *PWT7* in *P. oryzae.* The presence/absence of their functional/nonfunctional homologs was plotted on a phylogenetic tree constructed with RAxML-ng software, using the biallelic single nucleotide polymorphism sites in 3000 BUSCO genes. *PBY2* includes *PBY1* in MoE (EC-II).

Molecular mapping with the BC_1_F_1_ population narrowed down the target region to 3.5cM flanked by molecular markers, POTS50_2 and POTS65_1 (Fig. 3B), which corresponded to 580kb (Fig. 3C) in the Br48 whole genome sequence (GCA_036493215.1). A BAC library of Br48 was screened with molecular markers designed in this region, and four clones identified (Fig. 3C) were introduced into 200R45. BAC1, which turned 200R45 avirulent on R81, was digested with BbvCI, subcloned, and sequenced. Predicted genes encoding putative effector-like proteins were introduced into 200R45, and resulting transformants were sprayed on R81. Only one gene, ORF40, turned 200R45 avirulent on R81 (Fig. 3D), suggesting that ORF40 is *PBY2*. *PBY2* encoded a MAX (*Magnaporthe* AVRs and ToxB-like) effector with a signal peptide (Fig. 3E), and was located on chromosome 1 of Br48 (Fig. 3F). When *PBY2* of Br48 was disrupted (fig. S5), the strain (Br48ΔPBY2) lost the avirulence on R81 (Fig. 3G). Conversely, when *PBY2* was re-introduced into the Br48ΔPBY2 strain, transformants carrying a few copies of the transgene and expressing *PBY2* at the comparable level to Br48 (fig. S6) re-gained the avirulence on R81 (Fig. 3G). Interestingly, transformants expressing *PBY2* at a higher level (fig. S6) due to many copies of the transgene were avirulent not only on R81 but also on R74 (Fig. 3G). These results suggest that *Rmo2.c* in R74 is able to recognize *PBY2* to some extent although the degree of the recognition is not enough to express resistance to a wild isolate, Br48, which carries only one copy of *PBY2*.

MZ5-1-6 also carried a single copy of ORF40 (ORF40^MZ^). ORF40^MZ^ was identical to the intact ORF40 in Br48 (ORF40^Br^) at the nucleotide sequence level. ORF40^MZ^ and ORF40^Br^ segregated independently in the F_1_ population derived from MZ5-1-6 x Br48 (fig. S7) as they are located on different chromosomes (Fig. 3F). The presence/absence of the ORF40 signal derived from MZ5-1-6 co-segregated with the avirulence/virulence of the F_1_ cultures on R74 (fig. S7).

These results suggest that ORF40^MZ^ is *PBY1* conditioning the avirulence of MZ5-1-6 on R81 and R74 (*36*). When *PBY1* of MZ5-1-6 was disrupted (fig. S5), the strain (MZ5-1-6ΔPBY1) lost the avirulence on R81 and R74 (Fig. 3G). Conversely, when *PBY1* was re-introduced into the MZ5-1-6ΔPBY1 strain, transformants carrying a few copies of the transgene and expressing *PBY1* at the comparable level to MZ5-1-6 (fig. S6) gained the avirulence on R81 and R74 (Fig. 3G).

Furthermore, transformants expressing *PBY1* at a higher level (fig. S6) due to many copies of the transgene were avirulent not only on R81 and R74 but also on H4 (Fig. 3G). Taken together, we conclude that the three alleles at the *RMO2* locus (*Rmo2.a*, *Rmo2.c*, and *Rmo2.d*) are all able to recognize ORF40 (*PBY1* and *PBY2*) but in different intensity.

Distribution of *PWT7* and *PBY2* in *P. oryzae* was surveyed using isolates from various hosts. *PWT7* was confined to MoE (EC-II) and MoL (Fig. 3H). On the other hand, *PBY2* (including *PBY1*) was carried by all isolates of MoE, MoL, MoT, and those from *Eragrostis* (Fig. 3H), suggesting that *Rmo2.d* is widely effective against these three pathotypes and *Eragrostis* isolates.

### The role of the HMA domain in recognition of effectors

*Rmo2* in barley and *Rwt7* in wheat are located on terminal regions of the short arms of homoeologous chromosomes 7H and 7D, respectively (fig. S8). These regions in 7H and 7D also contain *Rpg1* (Fig. 1C) and G012400 (Fig. 2C), respectively. Comparative analyses of amino acid sequences suggested that *Rwt7* and G012400 are wheat orthologs of *Rmo2* and *Rpg1*, respectively (Fig. 4A). There is an additional TKP gene located in this region of 7D, *WTK4* found in *Aegilops tauschii* (fig. S8). *WTK4* has a PKD-KD architecture and conditions resistance to the wheat powdery mildew fungus (*38*). A comparative analysis of its amino acid sequence suggests that *WTK4* is homologous to *Rmo2* and *Rwt7* rather than *Rpg1* and G012400 (Fig. 4A).

**Fig. 4.**
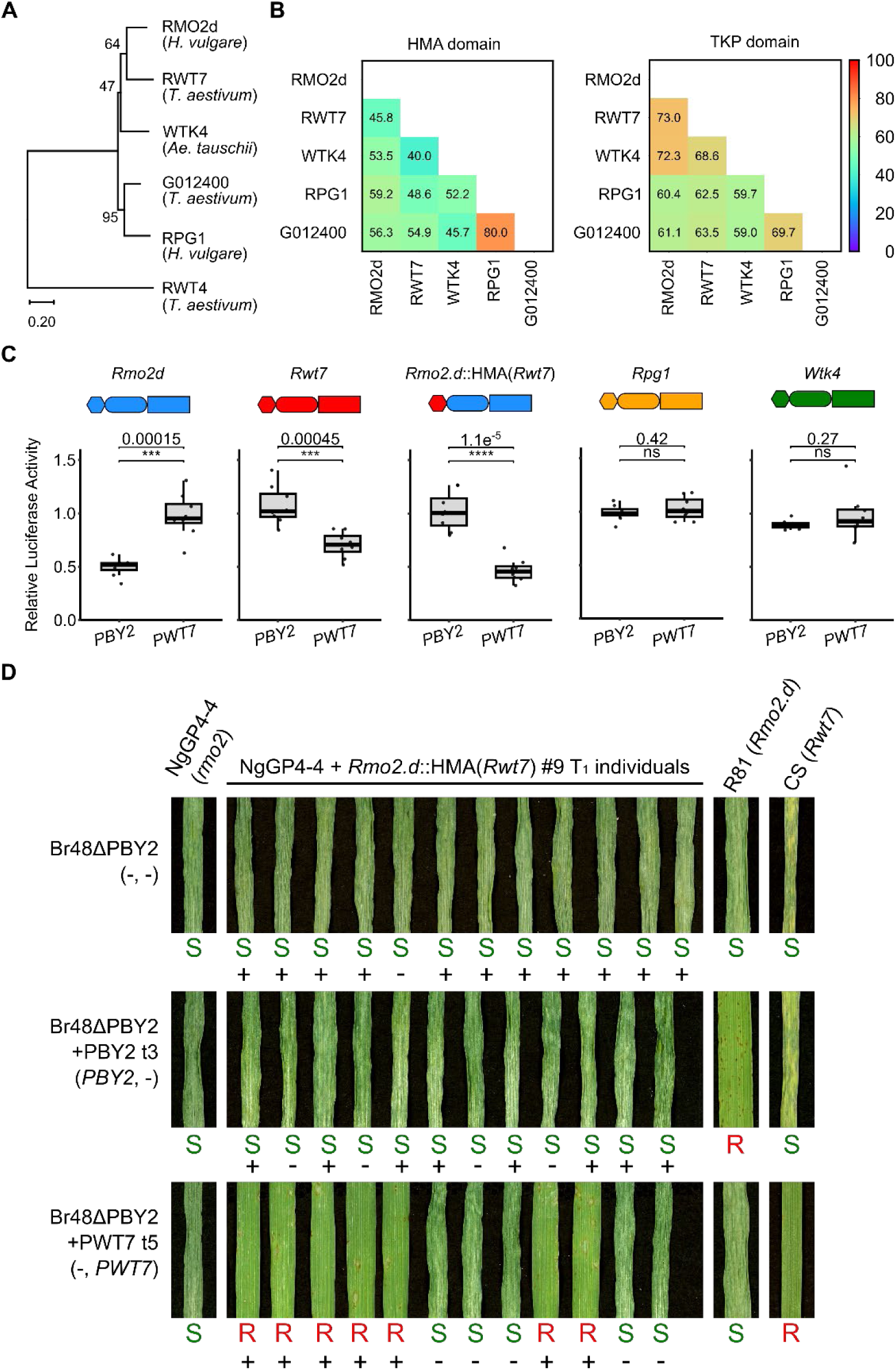
Role of HMA in effector recognition. (**A**) Maximum likelihood phylogenetic tree of representative HMA-integrated TKPs identified on homoeologous group 7 in barley (*Hordeum vulgare*), wheat (*Triticum aestivum*), and *Aegilops tauschii*. Bootstrap values from 1,000 replications are shown at nodes. RWT4, a TKP with a KD-PKD architecture, was used as an outgroup. (**B**) Heatmap showing amino acid sequence similarity (%) of domains of the TKPs. (**C**) Cell death assay with protoplasts. Protoplasts isolated from barley primary leaves were transfected with pAHC17-LUC containing a luciferase gene, pZH2Bik containing avirulence genes (*PBY2* or *PWT7* lacking signal peptides) or no insert (empty vector), and pZH2Bik containing *Rmo2.d*, *Rwt7*, *Rmo2.d*::HMA(*Rwt7*) (a construct encoding *Rmo2.d* protein kinases fused with Rwt7-HMA), *Rpg1*, or *WTK4*. Luciferase activity was determined 18-hours after transfection and represented as relative activities compared with those in samples with the empty vector. The experiments were repeated eight times. Results from the eight replicates were represented as a boxplot with original data points. Center lines show the medians; box limits indicate the 25th and 75th percentiles; whiskers extend to 1.5x the interquartile range from the 25th and 75th percentiles. Welch’s two-sided t-test was used for the comparisons of relative luciferase activity between the two groups. Triple and quadruple asterisks indicate significant differences at the 0.1% and 0.01% levels, respectively. NS, not significant. (**D**) Responses of barley transformants (T_1_) expressing *Rmo2.d*::HMA(*Rwt7*) to Br48ΔPBY2 and its transformant carrying either *PBY2* or *PWT7.* Genotypes of the fungal transformants are indicated in parentheses. Presence (+)/absence (-) of the transgene (*Rmo2.d*::HMA(*Rwt7*)) confirmed by PCR are shown below the panels. R, resistant; S, susceptible.

However, when comparing the HMA domains, the orthologous relationships are ambiguous except for the high homology between RPG1 and G012400 (Fig. 4B). This suggests that the diversification of HMA has played an important role in the expansion of recognized effectors. To investigate whether effector recognition is regulated by the HMA domains, we used a protoplast assay based on a luciferase reporter for cell viability. *Rmo2.d* and *Rwt7* specifically recognized *PBY2* and *PWT7*, respectively (Fig. 4C). Interestingly, when the HMA domain of *Rmo2.d* was replaced with that of *Rwt7*, the resulting construct, *Rmo2.d*::HMA(*Rwt7*), lost the ability to recognize *PBY2*, and gained the ability to recognize *PWT7* (Fig. 4C). *Rpg1* and *WTK4* did not recognize *PBY2* nor *PWT7*, suggesting that these genes may be adapted to recognize effectors from the stem rust fungus and the powdery mildew fungus, respectively.

To confirm the results obtained in the protoplast assay, *Rmo2.d*::HMA(*Rwt7*) was introduced into NgGP4-4. The T_1_ transformants were all susceptible to Br48ΔPBY2 and Br48ΔPBY2+PBY2 (a transformant of Br48ΔPBY2 carrying the *PBY2* transgene) (Fig. 4D). By contrast, against Br48ΔPBY2+PWT7 (a transformant of Br48ΔPBY2 carrying the *PWT7* transgene), resistant and susceptible individuals appeared in the T_1_ generation, and segregation of phenotype was perfectly correlated with presence/absence of the transgene (Fig. 4D). Taken together, we conclude that the HMA domain plays a critical role in recognizing target effectors.

### Distribution of *Rmo2* and *Rpg1* in the barley population

To survey the distribution of functional *Rmo2* in the barley population, transcripts were extracted from core barley accessions collected worldwide (fig. S9) and subjected to RT-PCR. Surprisingly, *Rmo2* alleles were amplified from all of the 113 accessions tested. This result indicates that *Rmo2* is expressed in all barley accessions. Furthermore, sequencing of these amplicons revealed that all of them maintained the *Rmo2* ORF.

On a dendrogram constructed using cDNA sequences of the kinase domains (PKD+KD), the *Rmo2* amplicons were divided into five groups, G1-G5 (fig. S9). This grouping was almost correlated with grouping of the HMA domain (figs. S9, S10). It should be noted that some of the G3 type of kinase domains were accompanied by two copies of HMA domains.

The kinase domains of *Rpg1* were mainly amplified from accessions carrying the G1 and G5 types of *Rmo2* (fig. S9). The *Rpg1* kinase domains from the G5 accessions were accompanied by the authentic *Rpg1* HMA. By contrast, those from the G1 accessions were accompanied by the HMA domain of the G5 type of *Rmo2* (fig. S9). This result suggests that this chimeric *Rpg1* gained its HMA from the G5 type of *Rmo2*.

## Discussion

The present study revealed that *Rmo2* in barley and *Rwt7* in wheat encode TKPs containing an HMA domain, and are orthologs located on syntenic regions of homoeologous chromosomes 7H and 7D, respectively. Interestingly, *Rmo2* was collocated with *Rpg1* controlling the resistance to the stem rust fungus, while *Rwt7* was collocated with *WTK4* controlling the resistance to the wheat powdery mildew fungus. *Rmo2* and *Rpg1* in barley are considered to have been derived from an ancient duplication that occurred before the differentiation of *Hordeum* and the *Triticum-Aegilops* complex because a corresponding paralogous pair of their orthologs, *Rwt7* and G012400, respectively, are present in wheat. Here, we designate the gene family comprising *Rmo2*, *Rwt7*, *Rpg1*, G012400, and *WTK4* as the “*Rpg1/Rmo2* family”. The evolutionary process of this family inferred from the present study is summarized in Fig. 5. The prototype of this family with a PKD-KD architecture emerged in the ancestral population of Triticeae, and evolved into a common ancestor of this family. The common ancestor duplicated, producing a paralog before the differentiation of *Hordeum* and the *Triticum-Aegilops* complex. This ancient paralog evolved into a pair of orthologs, *Rpg1* which recognizes the stem rust fungus and G012400 whose targets(s) remain unknown. The other paralog evolved into another pair of orthologs, *Rmo2* and *Rwt7*, both recognizing the blast fungus. In the *Triticum-Aegilops* complex *Rwt7* or its ancestor duplicated, giving rise to *WTK4* recognizing the powdery mildew fungus.

**Fig. 5.**
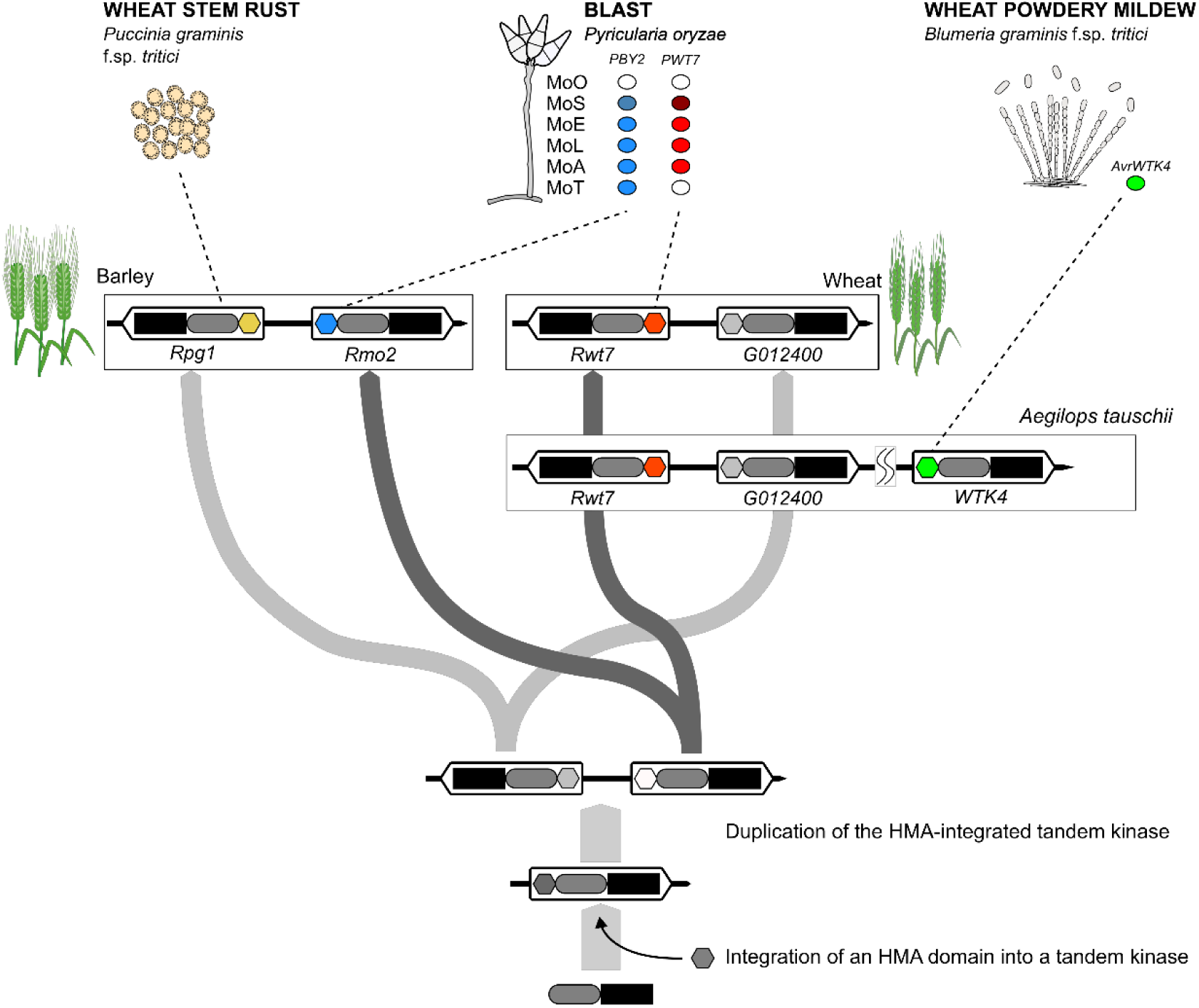
A model for the evolutionary trajectory of the *Rpg1/Rmo2* family encoding HMA-integrated TKPs. Hexagons, ellipses, and rectangles represent HMA, pseudokinase, and kinase domains, respectively. MoO, MoS, MoE, MoL, MoA, and MoT are pathotypes of *P. oryzae* (*<u>M</u>. <u>o</u>ryzae*) pathogenic on genus *<u>O</u>ryza*, *<u>S</u>etaria*, *<u>E</u>leusine*, *<u>L</u>olium, <u>A</u>vena,* and *<u>T</u>riticum*, respectively. Blue and red ovals represent functional *PBY2* and *PWT7*, respectively, carried by these pathotypes. Blue, red, and green hexagons represents HMA domains that recognize functional *PBY2*, *PWT7*, and *AvrWTK4*, respectively.

Expansion of target pathogens was also observed in another TKP gene family (*39–41*) and NLRs such as the MLA-like family (*9*; *42-45*). However, the molecular mechanisms underlying the target expansion have yet to be identified. Recently, Reveguk et al. (*19*) found that approximately half of TKPs detected in the plant kingdom contained integrated domains, and proposed that they could function as decoys for effectors. Bernasconi et al. (*46*) showed that the HMA domain of WTK4 directly interacts with the product of *AvrWTK4*. The present study suggested that the specific recognition of effectors or pathogens by the members of the *Rpg1/Rmo2* family is determined by the integrated HMA domains (Fig. 4). We propose that the diversification of the HMA domains enabled each member of the *Rpg1/Rmo2* family to shift their target effectors, thereby driving the expansion of target pathogens. It should be noted that all barley accessions tested (including the susceptible cultivar Ngt) carried and expressed an *Rmo2* allele with an undisrupted ORF (fig. S9). In addition, we showed that *Rmo2* can have a duplicated HMA or provided one of its HMA type to *Rpg1* (fig. S9). Throughout evolution, *RMO2* may have been an important, progenitrix locus that gave rise to paralogs, orthologs, and alleles effective against various effectors or pathogens.

Barley has been regarded as a universal suscept to *P. oryzae* because it is susceptible to various pathotypes (*35*). If an *Rmo2* construct recognizing both *PBY2* and *PWT7* is produced through molecular engineering, however, it may provide barley with robust, nonhost-like resistance against the *Eleusine* (EC-II), *Lolium*, and *Avena* pathotypes because these pathotypes carry both avirulence genes (Fig. 3H), and therefore, must mutate the two genes simultaneously to infect a barley transformant carrying the construct. A successful engineering of *Rmo2* for the dual recognition will be reported by Yu et al. (47).

## Acknowledgments

We thank M. J. Banfield, John Innes Centre, UK, for critical reading of, and valuable suggestions for the manuscript, and Reiko Kiuchi, Kobe, Japan, for financial supports. *Aegilops tauschii* accessions were provided by the National BioResource Project-Wheat with support in part by the National BioResource Project of the MEXT, Japan. This work was supported by the Joint Usage/Research Center, Institute of Plant Science and Resources, Okayama University.

## Author contributions

Conceptualizations: YT, SA

Methodology: YM, HH

Investigation: AGT, GSH, SK, TM, AH, DN, EK, MS, YT, MTI, NY, MS, HH

Funding acquisition: YT, SA

Project administration: YT, KS

Writing - original draft: YT, SA

Writing – review & editing: HH, KS, RT

## Data and materials availability

Sequence data for the genes described in the present study can be found in the GenBank/EMBL database under the accession numbers LC905591-LC905597. All plasmids, plant lines, and fungal strains generated in this work are available from the authors upon request.

## Supplementary Materials

Materials and Methods

Figs. S1 to S10

Tables S1 to S4

References (48–58)

## References and Notes

1. J. D. G. Jones, J. L. Dangl. The plant immune system. Nature 444,323–329 (2006).

2. J. Sánchez-Martin, B. Keller. NLR immune receptors and diverse types of non-NLR proteins control race-specific resistance in *Triticeae*. Curr. Opin. Plant Biol. 62, 102053 (2021).

3. G. Yu, O. Matny, S. Gourdoupis, N. Rayapuram, F. R. Aljedaani, Y. L. Wang, T. Nürnberger, R. Johnson, E. E. Crean, I. M.-L. Saur, C. Gardener, Y. Yue, N. Kangara, B. Steuernagel, S. Hayta, M. Smedley, W. Harwood, M. Patpour, S. Wu, J. Poland, J. D. G. Jones, T. L. Reuber, M. Ronen, A. Sharon, M. N. Rouse, S. Xu, K. Holušová, J. Bartoš, I. Molnár, M. Karafiátová, H. Hirt, I. Blilou, Ł. Jaremko, J. Doležel, B. J. Steffenson, B. B. H. Wulff. The wheat stem rust resistance gene *Sr43* encodes an unusual protein kinase. Nat. Genet. 55, 921–926 (2023).

4. J. D. G. Jones, B. J. Staskawicz, J. L. Dangl. The plant immune system: from discovery to deployment. Cell 187, 2095–2116 (2024).

5. J. L. Dangl, J. D. G. Jones. Plant pathogens and integrated defence responses to infection. Nature 411, 826–833 (2001).

6. R. A. L. van der Hoorn, S. Kamoun. From guard to decoy: a new model for perception of plant pathogen effectors. Plant Cell 20, 2009–2017 (2008).

7. L. Cadiou, F. Brunisholz, S. Cesari, T. Kroj. Molecular engineering of plant immune receptors for tailored crop disease resistance. Curr. Opin. Plant Biol. 74, 102381 (2023).

8. S. Cesari, M. Bernoux, P. Moncuquet, T. Kroj, P. N. Dodds. A novel conserved mechanisms for plant NLR protein pairs: the “integrated decoy” hypothesis. Front. Plant Sci. 5, 606 (2014).

9. S. G. Krattinger, B. Keller. Molecular genetics and evolution of disease resistance in cereals. New Phytol. 212, 320–332 (2016).

10. S. Cesari. Multiple strategies for pathogen perception by plant immune receptors. New Phytol. 219, 17–24 (2018).

11. Y. Okuyama, H. Kanzaki, A. Abe, K. Yoshida, M. Tamiru, H. Saitoh, T. Fujibe, H. Matsumura, M. Shenton, D. C. Galam, J. Undan, A. Ito, T. Sone, R. Terauchi. A multifaceted genomics approach allows the isolation of the rice *Pia*-blast resistance gene consisting of two adjacent NBS-LRR protein genes. Plant J. 66, 467–479 (2011).

12. I. Ashikawa, N. Hayashi, H. Yamane, H. Kanamori, J. Wu, T. Matsumoto, K. Ono, M. Yano. Two adjacent nucleotide-binding site-leucine-rich repeat class genes are required to confer *Pikm*-specific rice blast resistance. Genetics 180, 2267–2276 (2008).

13. H. Kanzaki, K. Yoshida, H. Saitoh, K. Fujisaki, A. Hirabuchi, L. Alaux, E. Fournier, D. Tharreau, R. Terauchi. Arms race co-evolution of *Magnaporthe oryzae AVR-Pik* and rice *Pik* genes driven by their physical interactions. Plant J. 72, 894–907 (2012).

14. S. Cesari, G. Thilliez, C. Ribot, V. Chalvon, C. Michel, A. Jauneau, S. Rivas, L. Alaux, H. Kanzaki, Y. Okuyama, J.-B. Morel, E. Fournier, D. Tharreau, R. Terauchi, T. Kroj. The rice resistance protein pair RGA4/RGA5 recognizes the *Magnaporthe oryzae* effectors AVR-Pia and AVR1-CO39 by direct binding. Plant Cell 25, 1463–1481 (2013).

15. A. Maqbool, H. Saitoh, M. Franceschetti, C. E. M. Stevenson, A. Uemura, H. Kanzaki, S. Kamoun, R. Terauchi, M. J. Banfield. Structural basis of pathogen recognition by an integrated HMA domain in a plant NLR immune receptor. eLife 4, e08709 (2015).

16. V. Klymiuk, G. Coaker, T. Fahima, C. Pozniak. Tandem protein kinases emerge as new regulators of plant immunity. Mol. Plant-Microbe Interact. 34, 1094–1102 (2021).

17. R. Brueggeman, N. Rostoks, D. Kudrna, A. Kilian, F. Han, J. Chen, A. Druka, B. Steffenson, Kleinhofs. The barley stem rust-resistance gene *Rpg1* is a novel disease-resistance gene with homology to receptor kinases. Proc. Natl. Acad. Sci. USA. 99, 9328–9333 (2002).

18. H. Horvath, N. Rostoks, R. Brueggeman, B. Steffenson, D. von Wettstein, A. Kleinhofs. Genetically engineered stem rust resistance in barley using the Rpg1 gene. Proc. Natl. Acad. Sci. USA. 100, 364–369 (2003).

19. T. Reveguk, A. Fatiukha, E. Potapenko, I. Reveguk, H. Sela, V. Klymiuk, Y. Li, C. Pozniak, T. Wicker, G. Coaker, T. Fahima,. Tandem kinase proteins across the plant kingdom. Nat. Genet. 57, 254–262 (2025).

20. E. Cavalet-Giorsa, A. González-Muñoz, N. Athiyannan, S. Holden, A. Salhi, C. Gardener, J. Quiroz-Chávez, S. M. Rustamova, A. F. Elkot, M. Patpour, A. Rasheed, L. Mao, E. S. Lagudah, S. K. Periyannan, A. Sharon, A. Himmelbach, J. C. Reif, M. Knauft, M. Mascher, N. Stein, N. Chayut, S. Ghosh, D. Perovic, A. Putra, A. B. Perera, C.-Y. Hu, G. Yu, H. I. Ahmed, K. D. Laquai, L. F. Rivera, R. Chen, Y. Wang, X. Gao, S. Liu, W. J. Raupp, E. L. Olson, J.-Y. Lee, P. Chhuneja, S. Kaur, P. Zhang, R. F. Park, Y. Ding, D.-C. Liu, W. Li, F. Y. Nasyrova, J. Dvorak, M. Abbasi, M. Li, N. Kumar, W. B. Meyer, W. H. P. Boshoff, B. J. Steffenson, O. Matny, P. K. Sharma, V. K. Tiwari, S. Grewal, C. J. Pozniak, H. S. Chawla, J. Ens, L. T. Dunning, J. A. Kolmer, G. R. Lazo, S. S. Xu, Y. Q. Gu, X. Xu, C. Uauy, M. Abrouk, S. Bougouffa, G. S. Brar, B. B. H. Wulff, S. G. Krattinger. Origin and evolution of the bread wheat D genome. Nature 633, 848–855 (2024).

21. R. Chen, J. Chen, O. R. Powell, M. A. Outram, T. Arndell, K. Gajendiran, Y. L. Wang, J. Lubega, Y. Xu, M. A. Ayliffe, C. Blundell, M. Figueroa, J. Sperschneider, T. Vanherche, K. Kanyuka, D. Tang, G. Zhong, C. Gardener, G. Yu, S. Gourdoupis, L. Jaremko, O. Matny, B. J. Steffenson, W. H. P. Boshoff, W. B. Meyer, S. T. Arold, P. N. Dodds, B. B. H. Wulff. A wheat tandem kinase activates an NLR to trigger immunity. Science 387 1402-1408 (2025).

22. P. Lu, G. Zhang, J. Li, Z. Gong, G. Wang, L. Dong, H. Zhang, G. Guo, M. Su, K. Wang, Y. Wang, K. Zhu, Q. Wu, Y. Chen, M. Li, B. Huang, B. Li, W. Li, L. Dong, Y. Hou, X. Cui, H. Fu, D. Qiu, C. Yuan, H. Li, J. Zhou, G.-Z Han, Y. Chen, Z. Liu. A wheat tandem kinase and NLR pair confers resistance to multiple fungal pathogens. Science 387, 1418–1424 (2025).

23. Y-C Sung, Y. Li, Z. Bernasconi, S. Baik, S. Asuke, B. Keller, T. Fahima, G. Coaker. Wheat tandem kinase RWT4 directly binds a fungal effector to activate defense. Nat. Genet. 57, 1238–1249 (2025).

24. H. Kato, M. Yamamoto, T. Yamaguchi-Ozaki, H. Kadouchi, Y. Iwamoto, H. Nakayashiki, Y. Tosa, S. Mayama, N. Mori. Pathogenicity, mating ability and DNA restriction fragment length polymorphisms of *Pyricularia* populations isolated from Gramineae, Bambusideae and Zingiberaceae Plants. J. Gen. Plant Pathol. 66, 30–47 (2000).

25. Y. Tosa, K. Hirata, H. Tamba, S. Nakagawa, I. Chuma, C. Isobe, J. Osue, A. S. Urashima, L. D. Don, M. Kusaba, H. Nakayashiki, A. Tanaka, T. Tani, N. Mori, S. Mayama. Genetic constitution and pathogenicity of *Lolium* isolates of *Magnaporthe oryzae* in comparison with host species-specific pathotypes of the blast fungus. Phytopathology 94, 454–462 (2004).

26. B. Valent, G. Cruppe, J. P. Stack, C. D. Cruz, M. L. Farman, P. A. Paul, G. L. Peterson, K. F. Pedley. Recovery plan for wheat blast caused by *Magnaporthe oryzae* pathotype *Triticum*. Plant Heal. Prog. 22, 182–212 (2021).

27. S. Asuke, M. Tanaka, F.S. Hyon, Y. Inoue, T. T. P. Vy, D. Niwamoto, H. Nakayashiki, Y. Tosa. Evolution of an *Eleusine*-specific subgroup of *Pyricularia oryzae* through a gain of an avirulence gene. Mol. Plant-Microbe Interact. 33, 153–165 (2020).

28. A. S. Urashima, S. Igarashi, H. Kato. Host range, mating type, and fertility of *Pyricularia grisea* from wheat in Brazil. Plant Dis. 77, 1211–1216 (1993).

29. M. Latorre, V. M. Were, A. J. Foster, T. Langner, A. Malmgren, A. Harant, S. Asuke, S. Reyes-Avila, D. R. Gupta, C. Jensen, W. Ma, N. U. Mahmud, M. S. Mehebub, R. M. Mulenga, A. N. M. Muzahid, S. K. Paul, S. M. F. Rabby, A. A. M. Rahat, L. Ryder, R.-K. Shrestha, S. Sichilima, D. M. Soanes, P. K. Singh, A. R. Bentley, D. G. O. Saunders, Y. Tosa, D. Croll, K. H. Lamour, T. Islam, B. Tembo, J. Win, N. J. Talbot, H. A. Burbano, S. Kamoun. Genomic surveillance uncovers a pandemic clonal lineage of the wheat blast fungus. PLoS Biol. 21, e3002052 (2023).

30. Y. Inoue, T. T. P. Vy, K. Yoshida, H. Asano, C. Mitsuoka, S. Asuke, V. L. Anh, C. J. R. Cumagun, I. Chuma, R. Terauchi, K. Kato, T. Mitchell, B. Valent, M. Farman, Y. Tosa. Evolution of the wheat blast fungus through functional losses in a host specificity determinant. Science 357, 80–83 (2017).

31. S. Asuke, S. Nishimi, Y. Tosa. At least five major genes are involved in the avirulence of an *Eleusine* isolate of *Pyricularia oryzae* on common wheat. Phytopathology 110, 465–471 (2020).

32. S. Asuke, N. J. Magculia, Y. Inoue, T. T. P. Vy, Y. Tosa. Correlation of genomic compartments with contrastive modes of functional losses of host specificity determinants during pathotype differentiation in *Pyricularia oryzae*. Mol. Plant-Microbe Interact. 34, 680–690 (2021).

33. S. Asuke, Y. Umehara, Y. Inoue, T. T. P. Vy, M. Iwakawa, Y. Matsuoka, K. Kato, Y. Tosa. Origin and dynamics of *Rwt6*, a wheat gene for resistance to nonadapted pathotypes of *Pyricularia oryzae*. Phytopathology 111, 2023–2029 (2021).

34. S. Asuke, A. Horie, K. Komatsu, R. Mori, T. T. P. Vy, Y. Inoue, Y. Jiang, Y. Tatematsu, M. Shimizu, Y. Tosa. Loss of *PWT7*, located on a supernumerary chromosome, is associated with parasitic specialization of *Pyricularia oryzae* on wheat. Mol. Plant-Microbe Interact. 36, 716–725 (2023).

35. G.-S. Hyon, N. T. T. Nga, I. Chuma, Y. Inoue, H. Asano, N. Murata, M. Kusaba, Y. Tosa. Characterization of interactions between barley and various host-specific subgroups of *Magnaporthe oryzae* and *M. grisea*. J. Gen. Plant Pathol. 78, 237–246 (2012).

36. N. T. T. Nga, Y. Inoue, I. Chuma, G-S. Hyon, K. Okada, T. T. P. Vy, M. Kusaba, Y. Tosa. Identification of a novel locus *Rmo2* conditioning resistance in barley to host-specific subgroups of *Magnaporthe oryzae*. Phytopathology 102, 674–682 (2012).

37. R. Brueggeman, T. Drader, A. Kleinhofs. The barley serin/threonine kinase gene *Rpg1* providing resistance to stem rust belongs to a gene family with five other members encoding kinase domains. Theor. Appl. Genet. 113, 1147–1158 (2006).

38. K. Gaurav, S. Arora, P. Silva, J. Sánchez-Martín, R. Horsnell, L. Gao, G. S. Brar, V. Widrig, W. J. Raupp, N. Singh, S. Wu, S. M. Kale, C. Chinoy, P. Nicholson, J. Quiroz-Chávez, J. Simmonds, S. Hayta, M. A. Smedley, W. Harwood, S. Pearce, D. Gilbert, N. Kangara, C. Gardener, M. Forner-Martínez, J. Liu, G. Yu, S. A. Boden, A. Pascucci, S. Ghosh, A. N. Hafeez, T. O’Hara, J. Waites, J. Cheema, B. Steuernagel, M. Patpour, A. F. Justesen, S. Liu, J. C. Rudd, R. Anvi, A. Sharon, B. Steiner, R. P. Kirana, H. Buerstmayr, A. A. Mehrabi, F. Y. Nasyrova, N. Chayut, O. Matny, B. J. Steffenson, N. Sandhu, P. Chhuneja, E. Lagudah, A. F. Elkot, S. Tyrrell, X. Bian, R. P. Davey, M. Simonsen, L. Schauser, V. K. Tiwari, H. R. Kutcher, P. Hucl, A. Li, D.-C. Liu, L. Mao, S. Xu, G. Brown-Guedira, J. Faris, J. Dvorak, M.-C. Luo, K. Krasileva, T. Lux, S. Artmeier, K. F. X. Mayer, C. Uauy, M. Mascher, A. R. Bentley, B. Keller, J. Poland, B. B. H. Wulff. Population genomic analysis of *Aegilops tauschii* identifies targets for bread wheat improvement. Nat. Biotechnol. 40, 422–431 (2022).

39. P. Lu, L. Guo, Z. Wang, B. Li, J. Li, Y. Li, D. Qiu, W. Shi, L. Yang, N. Wang, G. Guo, J. Xie, Q. Wu, Y. Chen, M. Li, H. Zhang, L. Dong, P. Zhang, K. Zhu, D. Yu, Y. Zhang, K. R. Deal, N. Huo, C. Liu, M.-C. Luo, J. Dvorak, Y. Q. Gu, H. Li, Z. Liu. A rare gain of function mutation in a wheat tandem kinase confers resistance to powdery mildew. Nat. Commun. 11, 680 (2020).

40. S. Arora, A. Steed, R. Goddard, K. Gaurav, T. O’Hara, A. Schoen, N. Rawat, A. F. Elkot, A. V. Korolev, C. Chinoy, M. H. Nicholson, S. Asuke, R. Antoniou-Kourounioti, B. Steuernagel, G. Yu, R. Awal, M. Forner-Martínez, L. Wingen, E. Baggs, J. Clarke, D. G. O. Saunders, K. V. Krasileva, Y. Tosa, J. D. G. Jones, V. K. Tiwari, B. B. H. Wulff, P. Nicholson. A wheat kinase and immune receptor form host-specificity barriers against the blast fungus. Nat. Plants 9, 385–392 (2023).

41. G. Yu, O, Matny, N. Champouret, B. Steuernagel, M. J. Moscou, I. Hernández-Pinzón, P. Green, S. Hayta, M. Smedley, W. Harwood, N. Kangara, Y. Yue, C. Gardener, M. J. Banfield, P. D. Olivera, C. Welchin, J. Simmons, E. Millet, A. Minz-Dub, Moshe Ronen, R. Avni, A. Sharon, M. Patpour, A. F. Justesen, M. Jayakodi, A. Himmelbach, N. Stein, S. Wu, J. Poland, J. Ens, C. Pozniak, M. Karafiátová, Istvan Molnár, J. Doležel, E. R. Ward, T. L. Reuber, J. D. G. Jones, M. Mascher, B. J. Steffenson, B. B. H. Wulff. *Aegilops sharonensis* genome-assisted identification of stem rust resistance gene *Sr62*. Nat. Commun. 13, 1607 (2022).

42. S. Periyannan, J. Moore, M. Ayliffe, U. Bansal, X. Wang, L. Huang, K. deal, M. Luo, X. Kong, H. Bariana, R. Mago, R. McIntosh, P. Dodds, J. Dvorak, E. Lagudah. The gene *Sr33*, an ortholog of barley *Mla* genes, encodes resistance to wheat stem rust race Ug99. Science 341, 786–788 (2013).

43. R. Mago, P. Zhang, S. Vautrin, H. Šimková, U. Bansal, M.-C. Luo, M. Rouse, H. Karaoglu, S. Periyannan, J. Kolmer, Y. Jin, M. A. Ayliffe, H. Bariana, R. F. Park, R. McIntosh, J. Doležel, H. Bergès, W. Spielmeyer, E. S. Lagudah, J. G. Ellis, P. N. Dodds. The wheat *Sr50* gene reveals rich diversity at a cereal disease resistance locus. Nat. Plants 1, 15186 (2015).

44. J. Bettgenhaeuser, I. Hernández-Pinzón, A. M. Dawson, M. Gardiner, P. Green, J. Taylor, M. Smoker, J. N. Ferguson, P. Emmrich, A. Hubbard, R. Bayles, R. Waugh, B. J. Steffenson, B. B. H. Wulff, A. Dreiseitl, E. R. Ward, M. J. Moscou. The barley immune receptor *Mla* recognizes multiple pathogens and contributes to host range dynamics. Nat. Commun. 12, 6915 (2021).

45. H. J. Brabham, D. G. D. L. Cruz, V. Were, M. Shimizu, H. Saitoh, I. Hernández-Pinzón, P. Green, J. Lorang, K. Fujisaki, K. Sato, I. Molnár, H. Šimková, J. Doležel, J. Russell, J. Taylor, M. Smoker, Y. K. Gupta, T. Wolpert, N. J. Talbot, R. Terauchi, M. J. Moscou. Barley MLA3 recognizes the host-specificity effector Pwl2 from *Magnaporthe oryzae*. Plant Cell 36, 447–470 (2024).

46. Z. Bernasconi, U. Stirnemann, Y. Xu, A. G. Herger, R. Chen, V. Widrig, M. Heuberger, M. Lettieri, M. C. Müller, B. B. H. Wulff, T. Wicker, B. Keller, J. Sánchez-Martín. An HMA-like integrated domain in the wheat tandem kinase WTK4 recognizes an RNase-like pathogen effector. bioRxiv. 10.1101/2025.08.26.672365 (2025).

47. D. S. Yu, R. Zdrzałek, E. Katayama, H. Akiyama, L. Daykin, N. J. Williams, I. Goodridge, S. Asuke, M. J. Banfield. Protein engineering of plant tandem kinase immune receptors expands effector recognition profiles. co-submitted (2025).

48. J. Murakami, Y. Tosa, T. Kataoka, R. Tomita, J. Kawasaki, I. Chuma, Y. Sesumi, M. Kusaba, H. Nakayashiki, S. Mayama. Analysis of host species specificity of *Magnaporthe grisea* toward wheat using a genetic cross between isolates from wheat and foxtail millet. Phytopathology 90, 1060–1067 (2000).

49. A. G. Tagle, I. Chuma, Y. Tosa. *Rmg7*, a new gene for resistance to *Triticum* isolates of *Pyricularia oryzae* identified in tetraploid wheat. Phytopathology 105, 495–499 (2015).

50. H. Hisano, K. Sato. Genomic regions responsible for amenability to Agrobacterium-mediated transformation in barley. Sci. Rep. 6, 37505 (2016).

51. Y. Tosa, J. Osue, Y. Eto, H. S. Oh, H. Nakayashiki, S. Mayama, S. A. Leong. Evolution of an avirulence gene *AVR1-CO39* concomitant with the evolution and differentiation of *Magnaporthe oryzae*. Mol. Plant-Microbe Interact. 18, 1148–1160 (2005).

52. I. M. L. Saur, S. Bauer, X. Lu, P. Schulze-Lefert. A cell death assay in barley and wheat protoplasts for identification and validation of matching pathogen AVR effector and plant NLR immune receptors. Plant methods 15, 118 (2019).

53. H. Hisano, B. Meints, M. J. Moscou, L. Cistue, B. Echávarri, K. Sato, P. M. Hayes. Selection of transformation-efficient barley genotypes based on *TFA* (transformation amenability) haplotype and higher resolution mapping of the *TFA* loci. Plant Cell Rep. 36, 611–620 (2017).

54. M. Shimizu, Y. Nakano, A. Hirabuchi, K. Yoshino, M. Kobayashi, K. Yamamoto, R. Terauchi, H. Saitoh. RNA-Seq of in planta-expressed *Magnaporthe oryzae* genes identifies MoSVP as a highly expressed gene required for pathogenicity at the initial stage of infection. Mol. Plant Pathol. 20, 1682–1695 (2019).

55. M. Shimizu, A. Hirabuchi, Y. Sugihara, A. Abe, T. Takeda, M. Kobayashi, Y. Hiraka, E. Kanzaki, K. Oikawa, H. Saitoh, T. Langner, M. J. Banfield, S. Kamoun, R. Terauchi. A genetically linked pair of NLR immune receptors shows contrasting patterns of evolution. Proc. Natl Acad. Sci. USA 119, e2116896119 (2022).

56. H. Chiapello, L. Mallet, C. Guérin, G. Aguileta, J. Amselem, T. Kroj, E. Ortega-Abboud, M.-H. Lebrun, B. Henrissat, A. Gendrault, F. Rodolphe, D. Tharreau, E. Fournier. Deciphering genome content and evolutionary relationships of isolates from the fungus *Magnaporthe oryzae* attacking different host plants. Genome Biol. Evol. 7, 2896–2912 (2015).

57. K, Yoshida, D. G. Saunders, C. Mitsuoka, S. Natsume, S. Kosugi, H. Saitoh, Y. Inoue, I. Chuma, Y. Tosa, L. M. Cano, S. Kamoun, R. Terauchi. Host specialization of the blast fungus *Magnaporthe oryzae* is associated with dynamic gain and loss of genes linked to transposable elements. BMC Genomics 17, 370 (2016).

58. Z. Peng, E. Oliveira-Garcia, G. Lin, Y. Hu, M. Dalby, P. Migeon, H. Tang, M. Farman, D. Cook, F. F. White, B. Valent, S. Liu. Effector gene reshuffling involves dispensable mini-chromosomes in the wheat blast fungus. PLoS Genet. 15, e1008272 (2019).

