## Supplementary material for "Evolution of HMA-integrated tandem kinases accompanied by expansion of target pathogens": figure S1-S10, table S1-S4

**The PDF file includes:**

Materials and Methods
Figs. S1 to S10
Tables S1 to S4

### Materials and Methods

#### Plant materials

*Hordeum vulgare* 'Nigrate' (Ngt), 'H.E.S.4' (H4), 'Morex' (Mrx), 'Russia 74' (R74), 'Golden Promise' (GP), and 'Russia 81' (R81) were used as representative cultivars for isolation of *Rmo2*. *Triticum aestivum* 'Transfed' (Tfed), 'Hope', 'Chinese Spring' (CS), and 'Norin 4' (N4) were used as representative cultivars for isolation of *Rwt7*. For distribution analysis of *Rmo2* and *Rpg1*, 112 lines were selected from 274 standard varieties of *H. vulgare* subsp. *vulgare* and 14 accessions of *H. vulgare* subsp. *spontaneum* (table S2) provided by the National BioResource Project Barley (NBRP; http://earth.nig.ac.jp/~dclust/cgi-bin/index.cgi).

#### Fungal materials

*Pyricularia oryzae* isolates used in the present study are listed in table S3. To produce the 86 random $BC_1F_1$ cultures for mapping of *PBY2*, 200R45 and 200R69, random $F_1$ cultures derived from Br48 (MoT isolate) × MZ5-1-6 (MoE isolate) (*31*) were backcrossed with Br48 on oatmeal agar media as described previously (48). All isolates/strains of *P. oryzae* have been maintained on sterilized barley seeds at Kobe University.

#### Infection assay

Wheat and barley seeds were pregerminated on moistened filter papers for 24 h, sown in vermiculite supplied with liquid fertilizer in a seeding case (5.5 × 15 × 10 cm), and grown at 22°C in a controlled-environment room with a 12-h photoperiod of fluorescent lighting for 8 days. Primary leaves of the 9-day-old seedlings were fixed onto a hard plastic board with rubber bands just before inoculation. Inocula ($1 \times 10^5$ conidia/ml) were prepared as described by Tagle et al. (49). The conidial suspension was sprayed onto primary leaves using an air compressor. The inoculated seedlings were incubated in dark and humid conditions at 22 to 25°C for 24 h, then returned to dry conditions with fluorescent lighting and incubated for an additional 3 to 5 days at 22 to 25°C. Four to six days after inoculation, symptoms (infection types) were evaluated on the basis of the size and color of lesions (*35*). The size was rated on six progressive grades from 0 to 5 as follows: 0, no visible infection; 1, pinhead spots; 2, small lesions (<1.5 mm); 3, scattered lesions of intermediate size (<3 mm); 4, large typical lesion; and 5, complete blighting of leaf blades. A disease score comprised a number denoting the lesion size and a letter indicating the lesion color: B for brown and G for green.

**Mapping of *Rmo2* and *Rwt7***

A total of 93 $F_{2:3}$ lines derived from Ngt x R81 were used for mapping of *Rmo2*. Twenty seeds were retrieved from each $F_{2:3}$ line and subjected to infection assay with GFSI1-7-2 for phenotyping. Another set of 20 seeds was retrieved from each $F_{2:3}$ line, sown in vermiculite, and grown at 22°C for 7 days. Seven-day-old primary leaves were bulked and used for DNA extraction with the CTAB method. Molecular markers were developed based on the barley genome sequence and a syntenic region in *Brachypodium distachyon* chromosome 1. Cleaved Amplified Polymorphic Sequence (CAPS) markers were designed using CAPS Designer (https://solgenomics.net/tools /caps_designer/caps_input.pl). Marker fragments were amplified from genomic DNA of the parental cultivars and the $F_{2:3}$ lines using Blend Taq and 2× Quick Taq HS DyeMix (TOYOBO, Osaka, Japan) following the manufacturer's instructions. Fragments amplified with primers for CAPS markers were digested with appropriate restriction enzymes supplied by Takara Bio (Kusatsu, Japan) or New England Biolabs Japan (Tokyo, Japan) (table S4). PCR products were electrophoresed in 0.7–2.0% agarose gels and stained with ethidium bromide for visualization. MAPMAKER/EXP version 3.0 was used for constructing a genetic map. The logarithm-of-odds (LOD) threshold for declaration of linkage was set at 4.0. Genetic distance was calculated with the Kosambi function. In fine mapping, genomic DNA was extracted each $F_3$ individual derived from segregating $F_{2:3}$ lines. Molecular markers were amplified from these DNA samples and detected as mentioned above.

Cloning of *Rwt7* was performed similarly. Rough mapping was conducted using 94 $F_{2:3}$ lines derived from CS x Tfed. The genomic sequence of the candidate region was retrieved from the CS genome v1.1 (https://plants.ensembl.org/Triticum_aestivum/Info/Index). Using primers designed on the basis of publicly available SNP information, corresponding portions of Tfed were amplified and sequenced by the Sanger sequencer. CAPS markers were subsequently developed using the identified sequence polymorphisms.

**Transformation of barley**

To produce a transformable barley line carrying the *rmo2* allele, $F_1$ plants derived from Ngt x GP were backcrossed with GP. From their $BC_1F_2$ progeny a transformable barley line (NgGP4-4) susceptible to GFSI1-7-2 and MZ5-1-6 was selected following the procedures described in fig. S3. The transformation vectors pBUH3-1037.d, pBUH3-H4a, pBUH3-Rwt7, and pBUH3-H7P2, which drive *Rmo2.d*, *Rmo2.a*, *Rwt7*, and *Rmo2.d*::HMA(*Rwt7*), respectively, under the control of the maize Ubiquitin 1 promoter, were introduced into NgGP4-4 via Agrobacterium-mediated transformation as described by Hisano and Sato (50). Insertions of transgenes were checked by PCR with the HPT primers (table S4).

**Transformation of *P. oryzae***

A BAC library of Br48 genomic DNA was constructed using the pcc1BAC/BamHI vector (Epicentre) and was screened with linked markers. A large plasmid DNA of BAC1, a clone containing the candidate region of *PBY2*, was extracted using NucleoBond Xtra BAC (Macherey-Nagel). BAC1 was partially digested with BbvCI, and fragments containing vector backbone was self-ligated, resulting in sub-clones to deduce the physical position of *PBY2*. Fragments containing *PBY2* candidates were amplified from genomic DNA of Br48, with KOD-Plus-Neo (Toyobo) and primers shown in table S4. Each fragment was cloned into pBluescript II SK(+). The BAC clone and these recombinant plasmids were introduced into 200R45 (*pby1*;*pby2*) via the polyethylene glycol–mediated co-transformation with pSH75 carrying a hygromycin B phosphotransferase gene as described by Tosa et al. (51). Introduced *PBY2* and *PBY1* were confirmed by PCR using the PBY2-ORF primers (table S4). *PBY2* and *PBY1* disruptants of Br48 and MZ5-1-6 were produced using the split marker method as described by Inoue et al. (30).

**Cell death assay with barley protoplasts**

Protoplast cell death assays were performed to examine the function of *Rmo2.d*, *Rwt7,* and their swapping construct. Total RNA was extracted from inoculated leaves of R81 and CS with Sepasol-RNA I Super G (Nacalai Tesque, Japan) and were treated with DNase I at 37°C for 20 min. Complementary DNA was synthesized using PrimeScript™ 1st strand cDNA Synthesis Kit (Takara, Japan). *Rmo2.d* and *Rwt7* were amplified from cDNA of R81 and CS, respectively. Similarly, *Rpg1* and *WTK4* were amplified from cDNA of Mrx and *Ae. tauschii* KU-2097, respectively. *Rmo2.d*::HMA(*Rwt7*) was constructed by replacing the HMA domain of *Rmo2.d* with *Rwt7*-HMA. The ORFs of *PBY2* and *PWT7* without signal peptides were amplified from genomic DNA of Br48 and Br58, respectively. Primers used for these PCR reactions are shown in table S4. All these fragments were cloned into the KpnI site of the pZH2Bik vector using In-Fusion Cloning Kit (Takara, Japan) so as to be driven by the rice ubiquitin promoter. Established plasmids were extracted by NucleoBond Xtra Maxi (Macherey-Nagel, Düren, Germany). Mesophyll protoplasts were prepared from eight-day-old primary leaves of NgGP4-4. Transfection assays with those plasmids were performed as described in Saur et al. (52). Briefly, plasmids containing AVR and resistance genes were introduced into the NgGP4-4 protoplasts with a plasmid containing the luciferase gene (pAHC17-LUC) via the polyethylene glycol treatment. After 18 hours incubation at 20°C in the dark, the protoplasts were lysed, and luciferase activity in the resulting cell extracts was measured for 1 second/well on the Tristar 3 luminometer mode (Berthold, Germany). The measured luminescence was normalized using the negative control in which the AVR gene was substituted with the empty pZH2Bik vector. This experiment was repeated eight times independently.

**Phylogenetic analysis of cDNA sequences of *Rmo2* and *Rpg1* in barley diversity panel**

*Rmo2* and *Rpg1* were amplified from cDNA of total RNA as described above and cloned into the EcoRV site of pBSIISK+. These fragments were sequenced with ABI capillary sequencers, and aligned with MAFFT (v7.520). Coding sequences were extracted from obtained sequences and concatenated. A maximum likelihood tree was constructed using MEGA X with 1,000 bootstrap replicates. Primers used in this section are listed in table S4.

| | HMA | | | | | | | | | | | | | | | | | | | | | | | | | | | | | | | |
|---|---|---|---|---|---|---|---|---|---|---|---|---|---|---|---|---|---|---|---|---|---|---|---|---|---|---|---|---|---|---|---|---|
| | 5 | 6 | 26 | 28 | 34 | 35 | 41 | 42 | 43 | 44 | 48 | 50 | 51 | 54 | 59 | 60 | 63 | 64 | 65 | 66 | 67 | 68 | 70 | 71 | 72 | 73 | 75 | 76 | 77 | 78 | 79 | 80 |
| R81 (*Rmo2.d*) | S | - | A | S | D | Q | K | H | G | K | V | V | V | V | R | M | L | F | P | N | A | Q | L | A | V | G | A | — | — | — | S | K |
| R74 (*Rmo2.c*) | . | K | . | . | . | . | . | . | . | . | . | . | . | . | . | . | . | . | . | . | . | . | . | . | . | . | . | K | E | R | . | . |
| H4 (*Rmo2.a*) | R | K | . | . | . | E | . | . | . | . | . | . | . | . | . | . | . | . | . | . | . | . | . | . | . | . | . | K | E | R | . | . |
| Ngt (*rmo2*) | C | K | S | A | E | K | R | D | D | M | I | L | F | I | K | L | V | S | T | H | V | Y | V | S | A | R | K | N | E | — | — | S |

| | PK1 | | | | | | | |
|---|---|---|---|---|---|---|---|---|
| | 86 | 87 | 93 | 94 | 101 | 103 | 187 | 202 |
| R81 (*Rmo2.d*) | E | F | N | D | I | R | F | A |
| R74 (*Rmo2.c*) | . | . | . | . | . | . | . | . |
| H4 (*Rmo2.a*) | . | . | . | . | . | . | . | T |
| Ngt (*rmo2*) | G | Y | D | F | V | H | L | . |

| | PK2 | | | | | | | | |
|---|---|---|---|---|---|---|---|---|---|
| | 476 | 494 | 546 | 586 | 636 | 647 | 667 | 682 | 690 |
| R81 (*Rmo2.d*) | R | A | C | P | S | K | V | R | S |
| R74 (*Rmo2.c*) | . | . | . | . | . | . | . | . | . |
| H4 (*Rmo2.a*) | . | . | . | Q | P | . | I | . | . |
| Ngt (*rmo2*) | G | G | G | Q | . | E | . | C | A |

**Fig. S1. Amino acid sequence alignment of representative *Rmo2* alleles.**

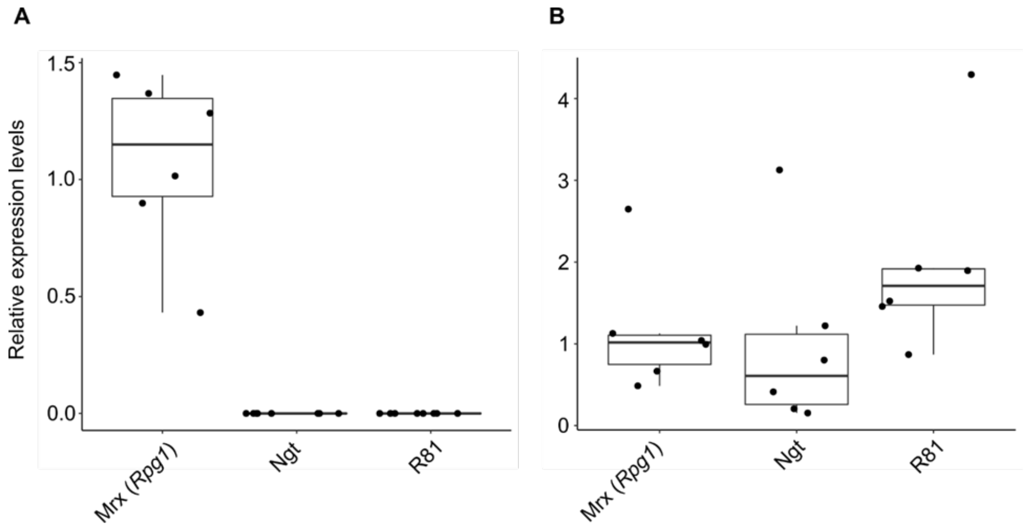


**Fig. S2. Expression of *Rpg1* (A) and *Rmo2* (B) in barley cultivars evaluated by quantitative PCR.**

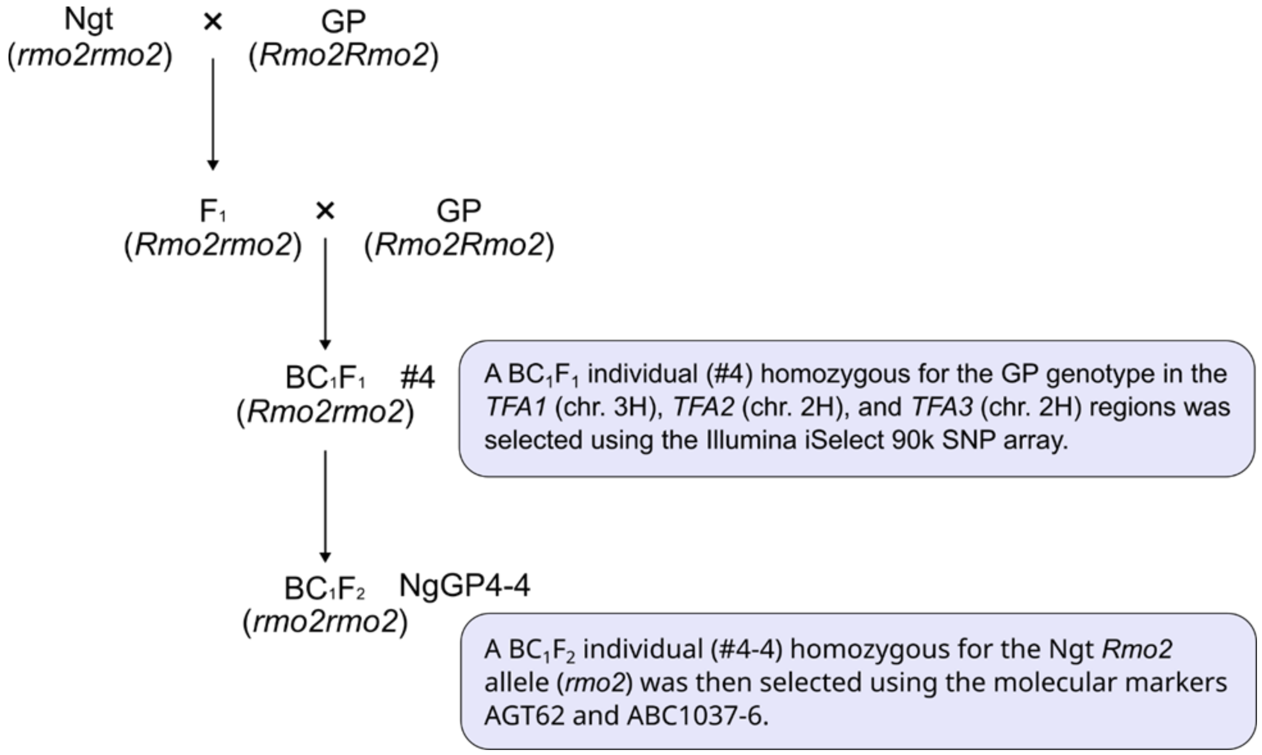


**Fig. S3. Pedigree of NgGP4-4, a transformable barley line carrying the *rmo2* allele.** *TFA1*, *TFA2*, and *TFA3* are three loci responsible for Transformation Amenability (*TFA*) to Agrobacterium-mediated transformation in barley (53).

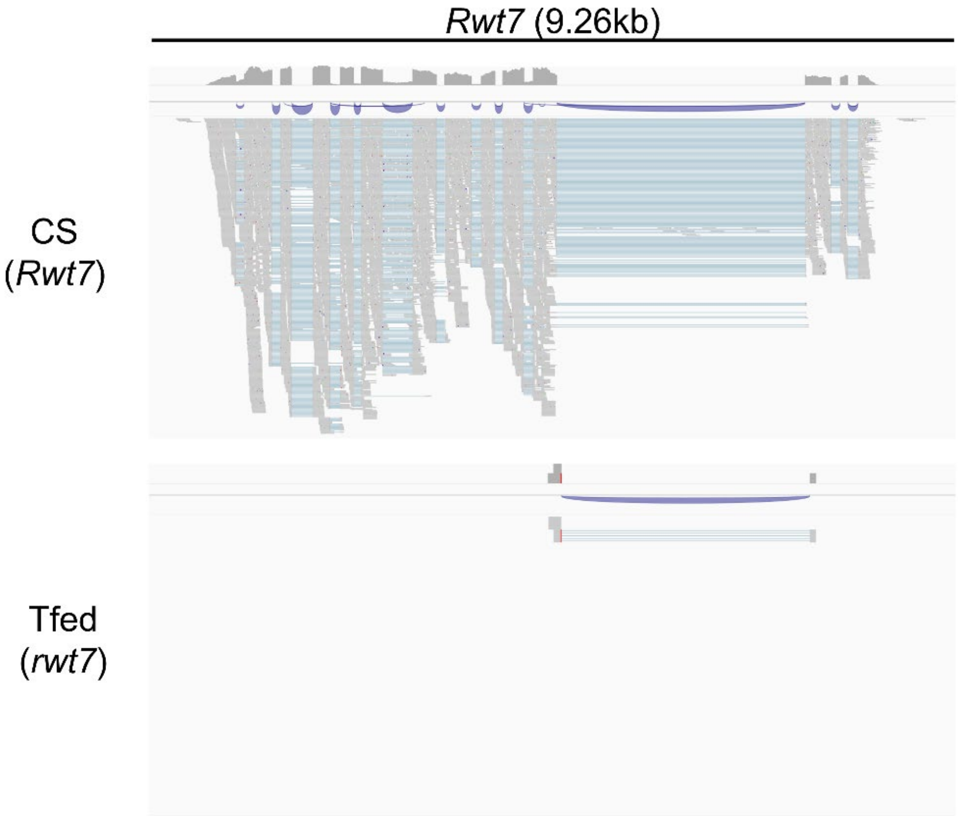


**Fig. S4. Alignment view of RNA short reads from CS and Tfed around G011800 (the *Rwt7* candidate) in the Chinese Spring v2.1 reference genome.**

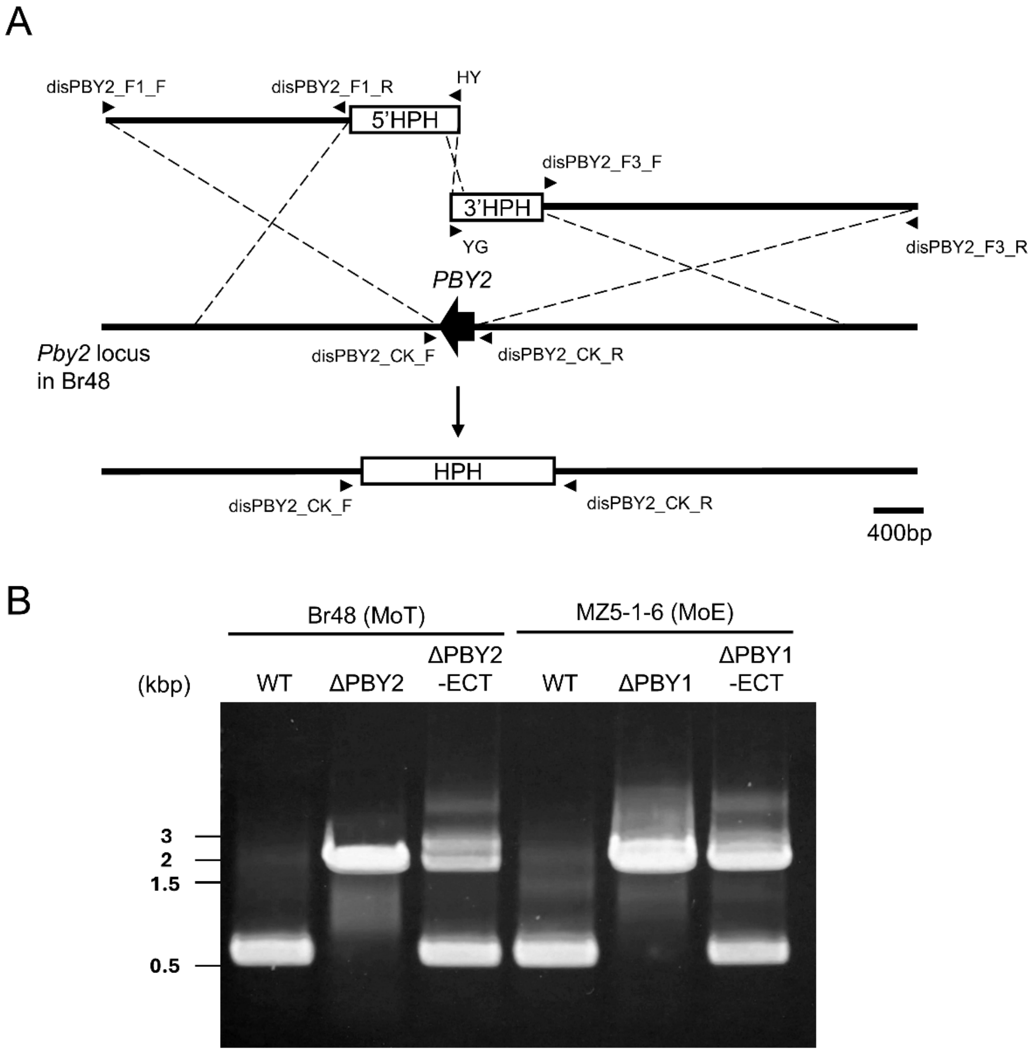


**Fig. S5. Targeted gene disruption of *PBY2* in Br48 and *PBY1* in MZ5-1-6.** (**A**) Split-marker strategy used for deletion of *PBY2* and *PBY1*. Regions indicated by crossover lines represent the sequences used for promoting homologous recombination. Primers used to amplify the targeted fragments are shown with arrowheads. Truncated (5′ HPH, 3′ HPH) or full-length HPH cassettes are depicted as rectangles. The overlapping region is 466 bp. (**B**) Verification of *PBY2* and *PBY1* deletion by PCR using the disPBY2_CK_F and disPBY2_CK_R primers. PCR products of 600 bp and 2,059 bp are expected from the wild-type and mutant genomic DNA, respectively. Marker sizes are shown on the left. WT, wild-type; ΔPBY2 and ΔPBY1, mutants with successful deletion of *PBY2* and *PBY1*, respectively; ΔPBY2-ECT and ΔPBY1-ECT, ectopic mutants.

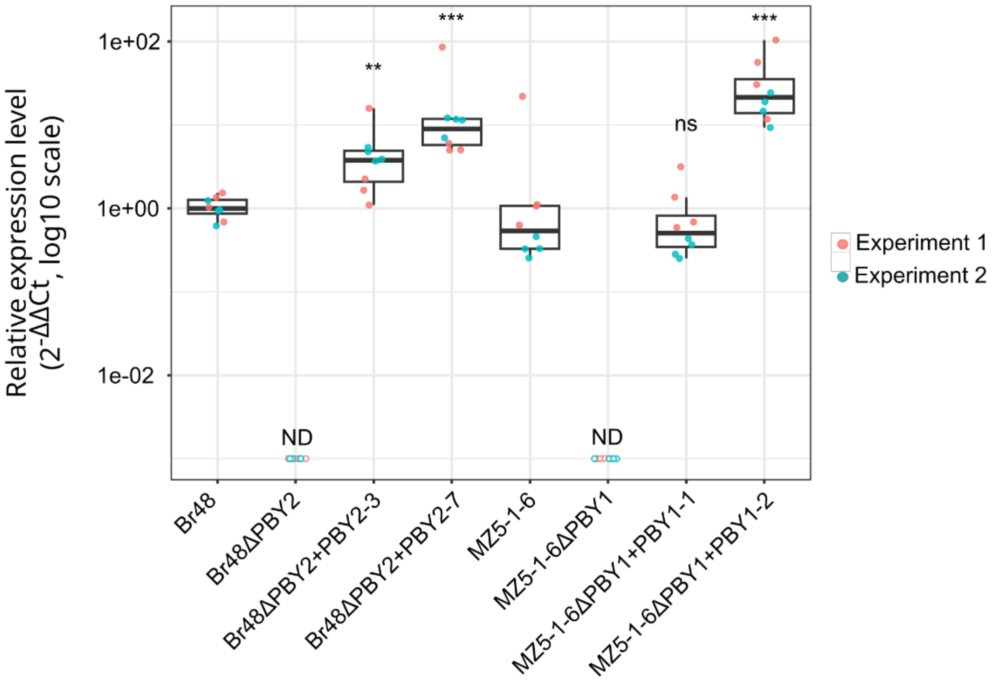


**Fig. S6. Relative expression levels of *PBY2* and *PBY1* in Br48 and MZ5-1-6, their disruptants, and transformants.** Transcript abundance was quantified using the $2^{-}\Delta\Delta$Ct method with Actin as the reference gene. Boxplots show biological replicates from two independent experiments. "ND" indicates reactions in which no detectable amplification occurred; ND values were excluded from statistical testing and plotted at a minimal value for visualization on a log scale. Significant differences among groups ($P < 0.001$) were detected by one-way ANOVA followed by Tukey's HSD test for pairwise comparisons. Adjusted significance levels (ns, **, ***) are shown above the plots.

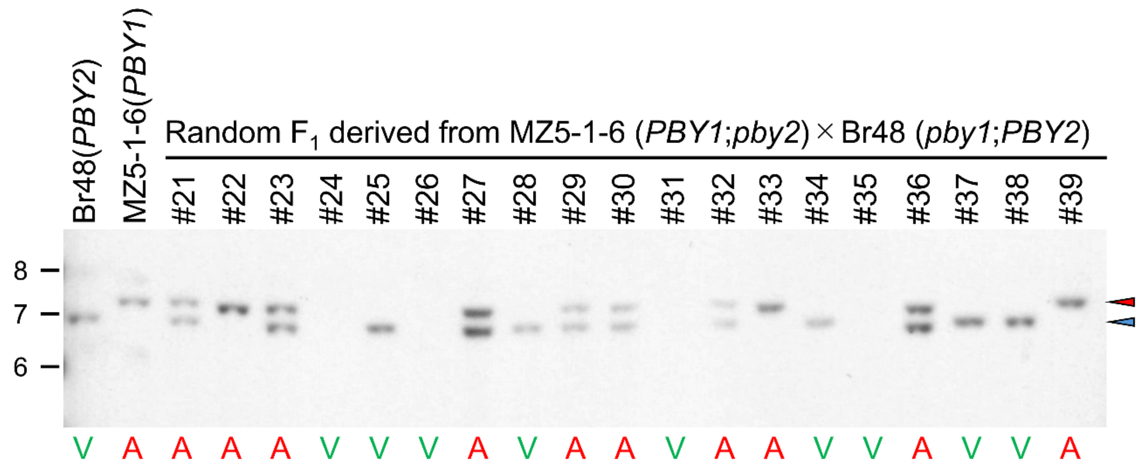


**Fig. S7. Segregation of *PBY2* and *PBY1* in $F_1$ cultures derived from MZ5-1-6 (*PBY1*;*pby2*) x Br48(*pby1*;*PBY2*).** Genomic DNA was digested with SalI, electrophoresed, blotted, and hybridized with pBluescript II SK(+) containing the fragment of *PBY2* open reading frame ($ORF40^{Br}$). Red and blue arrowheads indicate signals of *PBY1* (7.2 kb) and *PBY2* (6.8 kb), respectively. Phenotypes of these cultures on R74 are shown below the panel (A, avirulent; V, virulent).

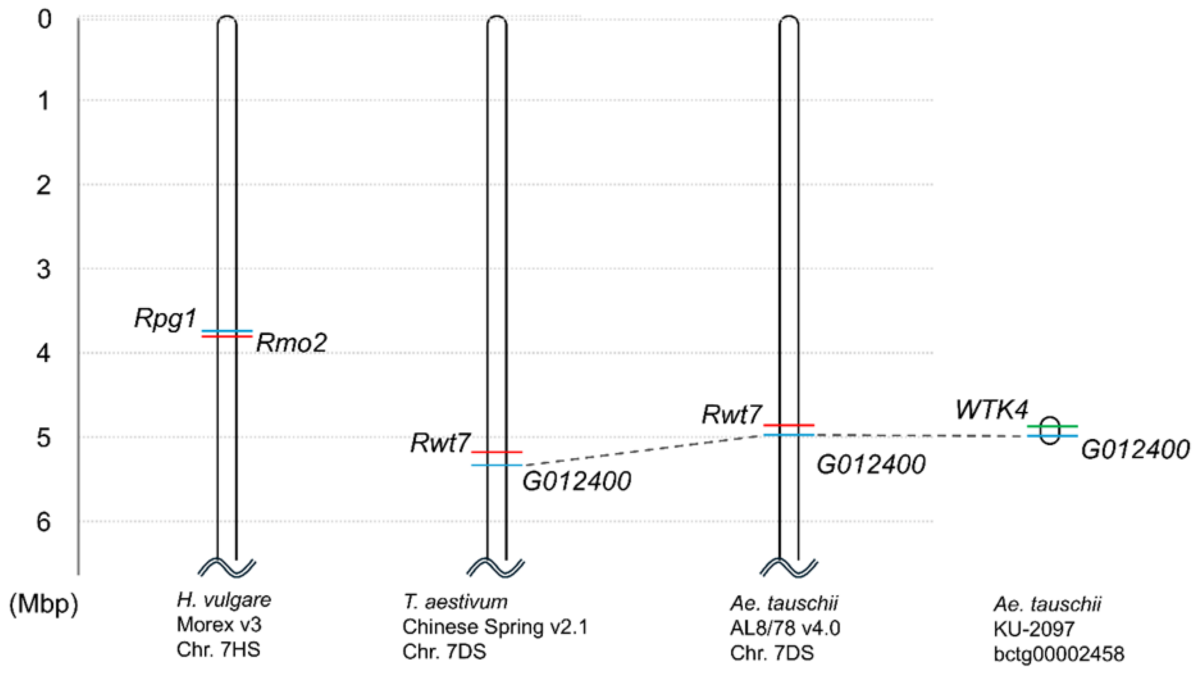


**Fig. S8. Physical maps of *Rmo2* and *Rwt7* in barley, wheat, and *Aegilops tauschii*.** KU-2097 is an *Ae. tauschii* accession carrying *WTK4* (*38*), which was identified by screening the National BioResource Project (NBRP) Komugi collection. Its whole genome was sequenced using the Oxford Nanopore Technologies PromethION sequencer. Bctg00002458 is a contig containing *WTK4* (322 kb in length).

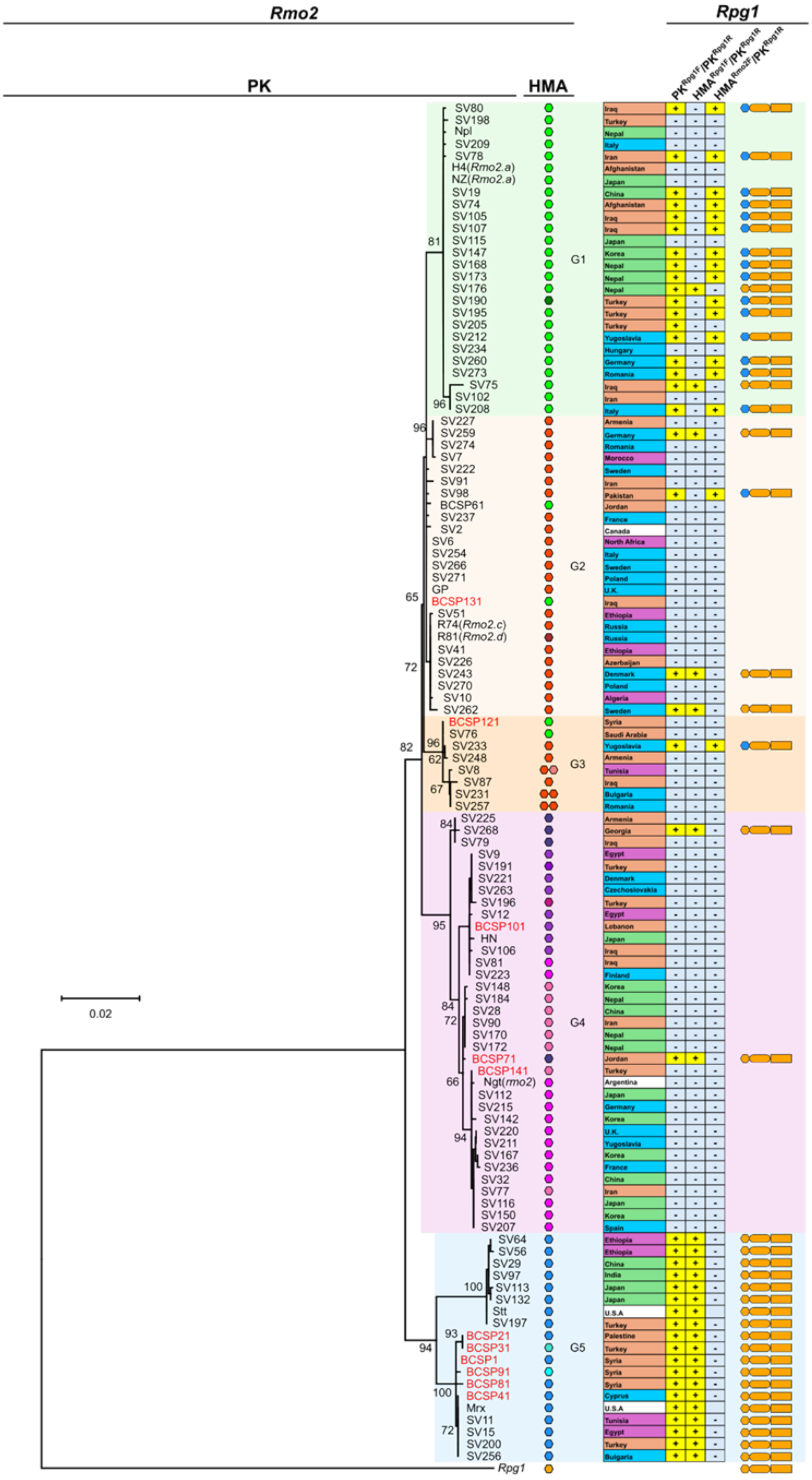
Rmo2
Rpg1
PK
HMA
PK^Rpg1F/PK^Rpg1R
HMA^Rpg1F/PK^Rpg1R
HMA^Rmo2F/PK^Rpg1R
G1
G2
G3
G4
G5
0.02
SV80 Iraq
SV198 Turkey
Npl Nepal
SV209 Italy
SV78 Iran
H4(Rmo2.a) Afghanistan
NZ(Rmo2.a) Japan
SV19 China
SV74 Afghanistan
SV105 Iraq
SV107 Iraq
SV115 Japan
SV147 Korea
SV168 Nepal
SV173 Nepal
SV176 Nepal
SV190 Turkey
SV195 Turkey
SV205 Turkey
SV212 Yugoslavia
SV234 Hungary
SV260 Germany
SV273 Romania
SV75 Iraq
SV102 Iran
SV208 Italy
SV227 Armenia
SV259 Germany
SV274 Romania
SV7 Morocco
SV222 Sweden
SV91 Iran
SV98 Pakistan
BCSP61 Jordan
SV237 France
SV2 Canada
SV6 North Africa
SV254 Italy
SV266 Sweden
SV271 Poland
GP U.K.
BCSP131 Iraq
SV51 Ethiopia
R74(Rmo2.c) Russia
R81(Rmo2.d) Russia
SV41 Ethiopia
SV226 Azerbaijan
SV243 Denmark
SV270 Poland
SV10 Algeria
SV262 Sweden
BCSP121 Syria
SV76 Saudi Arabia
SV233 Yugoslavia
SV248 Armenia
SV8 Tunisia
SV87 Iraq
SV231 Bulgaria
SV257 Romania
SV225 Armenia
SV268 Georgia
SV79 Iraq
SV9 Egypt
SV191 Turkey
SV221 Denmark
SV263 Czechoslovakia
SV196 Turkey
SV12 Egypt
BCSP101 Lebanon
HN Japan
SV106 Iraq
SV81 Iraq
SV223 Finland
SV148 Korea
SV184 Nepal
SV28 China
SV90 Iran
SV170 Nepal
SV172 Nepal
BCSP71 Jordan
BCSP141 Turkey
Ngt(rmo2) Argentina
SV112 Japan
SV215 Germany
SV142 Korea
SV220 U.K.
SV211 Yugoslavia
SV167 Korea
SV236 France
SV32 China
SV77 Iran
SV116 Japan
SV150 Korea
SV207 Spain
SV64 Ethiopia
SV56 Ethiopia
SV29 China
SV97 India
SV113 Japan
SV132 Japan
Stt U.S.A
SV197 Turkey
BCSP21 Palestine
BCSP31 Turkey
BCSP1 Syria
BCSP91 Syria
BCSP81 Syria
BCSP41 Cyprus
Mrx U.S.A
SV11 Tunisia
SV15 Egypt
SV200 Turkey
SV256 Bulgaria
Rpg1

**Fig. S9. Diversity of *Rmo2* and *Rpg1* in barley.** A maximum-likelihood phylogenetic tree was constructed using *Rmo2* cDNA sequences encoding kinase domains. Their HMA domains are shown by hexagon symbols which were color-coded according to the phylogenetic analyses of the HMA domain alone (Fig. S10). The kinase domains of *Rpg1* (in cv. Morex) were used as an outgroup. Numbers at the nodes represent bootstrap values (100 replicates). Accessions shown in red and black are *H. vulgare* ssp. *spontaneum* and *H. vulgare* ssp. *vulgare*, respectively. Their geographical origin (countries) is grouped into five regions: Africa (purple), Europe (blue), Middle East (orange), Asia (green), and North/South America (white). Shown to the right of the country's names are presence (+) /absence (-) of PCR products with forward/reverse primers, $PK^{Rpg1F}/PK^{Rpg1R}$, $HMA^{Rpg1F}/PK^{Rpg1R}$, and $HMA^{Rmo2F}/PK^{Rpg1R}$. $PK^{Rpg1F}$ and $PK^{Rpg1R}$ are designed for amplification of *Rpg1* kinase domains while $HMA^{Rpg1F}$ and $HMA^{Rmo2F}$ are forward primers designed for amplification of the entire genes (including the HMA-coding region ) of *Rpg1* and *Rmo2*, respectively.

**A**

| Representative cultivar | Numer of accessions | 1–41 |
|---|---|---|
| R81 (*Rmo2.d*) | 1 | MAAAS-KMVLKAELKDDKQKVKAVKALSVLHGIDQISVDMK |
| R74 (*Rmo2.c*) | 25 | .....K................................... |
| H4 (*Rmo2.a*) | 28 | ....RK............................E...... |
| SV190 | 1 | ....RK............................E...... |
| BCSP71 | 1 | ....RK...................S.A.....EK.N...R |
| SV268 | 3 | ....RK...................S.A.....EK.N...R |
| BCSP141 | 8 | ....CK...................S.A.....EK.....R |
| BCSP101 | 8 | ....CK...................S.A.....EK.....R |
| Ngt | 14 | ....CK...................S.A.....EK.....R |
| SV196 | 1 | ....CK...................S.A.....EK.....R |
| Mrx | 17 | ....RK.I.V.V.....S.CR..L....A.R..HV..ANPR |
| BCSP91 | 1 | ....RK.I.V.V....ES.CR..L....A.R..HV..ANPR |
| BCSP31 | 1 | .T..RK.I.V.V.....S.CR..L....A.R..HV..ANPR |

| Representative cultivar | 42–82 |
|---|---|
| R81 (*Rmo2.d*) | HGKITVVGVVDPVDVVARMR-KLFPNAQILAVGPA---SKY |
| R74 (*Rmo2.c*) | ....................-..............KER... |
| H4 (*Rmo2.a*) | ....................-..............KER... |
| SV190 | ..........Y.........-..............KER... |
| BCSP71 | DDM...I.LF..I....KL.-.VSTHVY.VSIP.ENE--S. |
| SV268 | DDM...I.LF.HI....KL.-.VSTHVY.IS.R.ENE--S. |
| BCSP141 | DDM...I.LF..I....KL.-.VSTHVY.VSAR.ENE--S. |
| BCSP101 | DDM...I.LF..I....KL.-.VSTHVY.VS.R.ENE--L. |
| Ngt | DDM...I.LF..I....KL.-.VSTHVY.VSAR.KNE--S. |
| SV196 | DDM...I.LF..I....KL.-.VSTH.Y.VS.R.ENE--L. |
| Mrx | ..N.....E.N.E..L..LQK........V....VME--TN |
| BCSP91 | ..N.....E.N.E..L..LQK........V.....KE--TN |
| BCSP31 | ..N.....E.N.E..L..LQK........V....VME--TN |

**B**

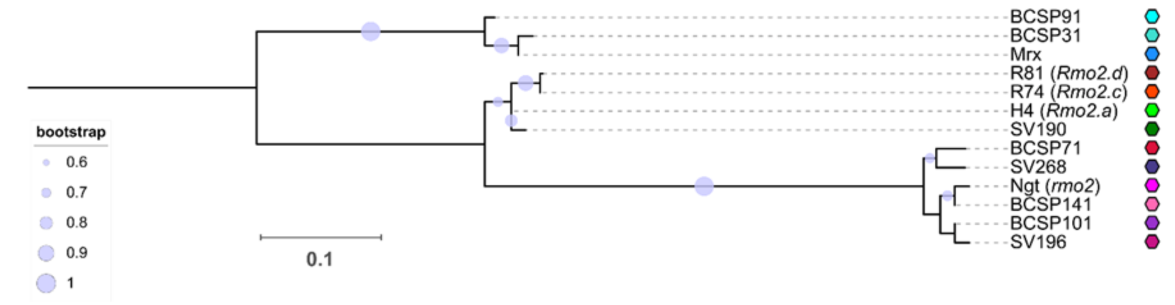


**Fig. S10. Diversity of HMA domains integrated into *Rmo2* in barley.** (**A**) Alignment of HMA domains found in *Rmo2* alleles. (**B**) A maximum likelihood tree of HMA domains of *Rmo2*. The dots on the branches were inferred from 1000 bootstrap replicates.

**Table S1. Segregation of reactions against Br48 + PWT7 in $F_2$ populations derived from crosses between wheat cultivars.**

| | Number of $F_2$ seedlings | | | | | | | | | | | |
|---|---|---|---|---|---|---|---|---|---|---|---|---|
| | Infection type | | | | | | | | | | | |
| Cross[a] | 0 | 1B | 2B | 3B | 4B | 4G | 5G | Total | R[b] | S[b] | $\chi^2$ (3:1) | P |
| CS x Tfed | 20 | 80 | 15 | 5 | 0 | 5 | 30 | 155 | 120 | 35 | 0.48 | 0.49 |
| Tfed x Hope | 13 | 74 | 23 | 5 | 0 | 6 | 33 | 154 | 115 | 39 | 0.01 | 0.93 |
| N4 x Tfed | 8 | 77 | 10 | 6 | 0 | 7 | 27 | 159 | 122 | 37 | 0.25 | 0.62 |
| CS x Hope | 11 | 266 | 17 | 0 | 0 | 0 | 0 | 294 | 294 | 0 | - | - |
| N4 x CS | 194 | 394 | 12 | 0 | 0 | 0 | 0 | 600 | 600 | 0 | - | - |
| N4 x Hope | 6 | 262 | 29 | 2 | 0 | 0 | 0 | 299 | 299 | 0 | - | - |

[a] CS, 'Chinese Spring'; Tfed, 'Transfed'; N4, 'Norin 4'

[b] R, resistant (0 to 4B); S, susceptible (4G to 5G)

**Table S2. Barley accessions used in the phylogenetic analysis**

| Species | Cultivar | Code | Country |
|---|---|---|---|
| *Hordeum vulgare* ssp. *vulgare* | Sanalta (CI 6087) | SV2 | Canada |
| | Libia | SV6 | North Africa |
| | Morocco | SV8 | Tunisia |
| | Giza 117 | SV9 | Egypt |
| | Macha | SV10 | Algeria |
| | Tunis (CI 1383) | SV11 | Tunisia |
| | Giza 68 | SV12 | Egypt |
| | Suez (84) | SV15 | Egypt |
| | Chiaochuang 1 | SV19 | China |
| | Paoanchen 1 | SV28 | China |
| | Shanghai 1 | SV29 | China |
| | Tinghsien | SV30 | China |
| | Takungkuan | SV32 | China |
| | Molale 1 (2-8-4a) | SV41 | Ethiopia |
| | Addis Ababa 3 (12-24-9) | SV51 | Ethiopia |
| | Adi Abun 1 (1-28-2) | SV56 | Ethiopia |
| | Sheki 2 (260-2) | SV64 | Ethiopia |
| | Kabul 1 | SV74 | Afghanistan |
| | Iraq Black Barley | SV75 | Iraq |
| | Aleppo 1 (438) | SV76 | Syria |
| | Esfahan 1 (152 III) | SV77 | Iran |
| | Gorgan 1 (225A I) | SV78 | Iran |
| | Khanaqin 2 (KUH 5305) | SV79 | Iraq |
| | Khanaqin 5 (KUH 5308) | SV80 | Iraq |
| | Arbat 1 (KUH 5317) | SV81 | Iraq |
| | Iraq Barley 1 | SV87 | Iraq |
| | Damaneh 1 (165d) | SV90 | Iran |
| | Ghazvin 1 (184) | SV91 | Iran |
| | Karad | SV97 | India |
| | Quetta 1 (14-1) | SV98 | Pakistan |

| | | |
|---|---|---|
| Ardabil 1 (336 II) | SV102 | Iran |
| Khanaqin 4 (KUH 5307) | SV105 | Iraq |
| Sulaymaniyah 2 (KUH 5322) | SV106 | Iraq |
| Sinjar 1 (KUH 5337) | SV107 | Iraq |
| Yahazu | SV112 | Japan |
| Shishikui Zairai | SV113 | Japan |
| Shimabara | SV115 | Japan |
| Sazanshu | SV116 | Japan |
| Wase Hadaka | SV132 | Japan |
| Waegwan Covered 1 | SV142 | Korea |
| Hamjong Covered 1 | SV147 | Korea |
| Hongweon Inuno-o | SV148 | Korea |
| Gwangju Covered 4 | SV150 | Korea |
| Hongcheon Covered 1 | SV167 | Korea |
| Sama 1 (1385) | SV168 | Nepal |
| Pisang 1 (1427) | SV170 | Nepal |
| Birkna Camp 1 (1487) | SV172 | Nepal |
| Kagbeni 1 (1616) | SV173 | Nepal |
| Gho 1 (1392) | SV176 | Nepal |
| Kakani Bangalow 3 (1477) | SV184 | Nepal |
| Turkey 31 | SV190 | Turkey |
| Turkey 61 | SV191 | Turkey |
| Turkey 41 | SV195 | Turkey |
| Turkey 71 | SV196 | Turkey |
| Turkey 101 | SV197 | Turkey |
| Rerhauli 1 (454) | SV198 | Turkey |
| Ayas (1309) | SV200 | Turkey |
| Istanbul (1024) | SV205 | Turkey |
| Samaria 4 Zeilige | SV207 | Spain |
| Bolognes | SV208 | Italy |
| Orayio | SV209 | Italy |
| Kolnozni | SV211 | Yugoslavia |
| Darmatishe | SV212 | Yugoslavia |
| Kleinwanz | SV215 | Germany |

| | | |
|---|---|---|
| Archer | SV220 | United Kingdom |
| Maja | SV221 | Denmark |
| Erhart Frederickson | SV222 | Sweden |
| Tammi | SV223 | Finland |
| Erevan 1 (Cauc.15) | SV225 | Armenia |
| Shemakha 1 (Cauc.37) | SV226 | Azerbaijan |
| Erevan 6 (Cauc.40) | SV227 | Armenia |
| Caveda | SV231 | Bulgaria |
| Urania | SV233 | Yugoslavia |
| Hungarian | SV234 | Hungary |
| Albert | SV236 | France |
| Cygne | SV237 | France |
| Opal | SV243 | Denmark |
| Geghard 1 (Cauc.44) | SV248 | Armenia |
| Tripoli | SV254 | Italy |
| Bulgarian 347 | SV256 | Bulgaria |
| Moldavia | SV257 | Rumania |
| Hanna | SV259 | Germany |
| Bavarian | SV260 | Germany |
| Tivannes | SV262 | Switzerland |
| Prokupkuy nahy | SV263 | Czechoslovakia |
| Ymer | SV266 | Sweden |
| Tibilisi 1 (Cauc.20) | SV268 | Georgia |
| PLD 49 | SV270 | Poland |
| PLD 83c | SV271 | Poland |
| KUH 836 | SV273 | Rumania |
| KUH 842 | SV274 | Rumania |
| Russia 81 | R81 | Russia |
| Russia 74 | R74 | Russia |
| H.E.S.4 | H4 | Afghanistan |
| Nakaizumi Zairai | NZ | Japan |
| Nigrate | Ngt | United State of America |
| Golden Promise | GP | United Kingdom |
| Morex | Mrx | United State of America |

| | | | |
|---|---|---|---|
| *H. vulgare* ssp. *spontaneum* | IG 38610 | BCSP1 | Syria |
| | IG 38661 | BCSP11 | Iran |
| | IG 38912 | BCSP21 | State of Palestine |
| | IG 39837 | BCSP31 | Turkey |
| | IG 39886 | BCSP41 | Cyprus |
| | IG 39933 | BCSP51 | Libya |
| | IG 40021 | BCSP61 | Jordan |
| | IG 40057 | BCSP71 | Jordan |
| | IG 40097 | BCSP81 | Syria |
| | IG 40194 | BCSP101 | Libya |
| | IG 107424 | BCSP111 | Iraq |
| | IG 110798 | BCSP121 | Syria |
| | IG 112797 | BCSP131 | Iraq |
| | IG 116121 | BCSP141 | Turkey |

**Table S3.** ***Pyricularia oryzae*** **isolates/strains used in this study**

| Isolate | Host | Locality | Year | Reference |
|---|---|---|---|---|
| Ina168 | *Oryza sativa* | Japan | 1958 | (*54*) |
| Ken53-33 | *O. sativa* | Japan | 1953 | (*34*) |
| PO12-7301-2 | *O. sativa* | Indonesia | 1973 | (*34*) |
| PH297 | *O. sativa* | Philippines | 2012 | (*34*) |
| 2012-01 | *O. sativa* | Japan | 1976 | (*55*) |
| Ao92-06-2 | *O. sativa* | Japan | 1992 | (*55*) |
| Ina87T-56A | *O. sativa* | Japan | 1987 | (*55*) |
| 85-141 | *O. sativa* | Japan | 1985 | (*55*) |
| SL91-48D | *O. sativa* | Japan | 1991 | (*55*) |
| 2403-1 | *O. sativa* | Japan | 1976 | (*55*) |
| Br15 | *O. sativa* | Brazil | 1990 | This study |
| VHG4.5 | *O. sativa* | Vietnam | 1996 | This study |
| H98-315-1 | *O. sativa* | Japan | 1998 | (*55*) |
| 0423-1 | *O. sativa* | Japan | 1976 | (*55*) |
| Ina85-182 | *O. sativa* | Japan | 1985 | (*55*) |
| Guy11 | *O. sativa* | French Guyana | 1978 | (*56*) |
| GFSI1-7-2 | *Setaria italica* | Japan | 1977 | (*34*) |
| NRSI2-2-2 | *S. italica* | Japan | 1977 | (*34*) |
| NRSI3-1-1 | *S. italica* | Japan | 1977 | (*34)* |
| NNSI3-2-1 | *S. italica* | Japan | 1984 | (*34)* |
| IN77-16-1-1 | *S. italica* | India | 1977 | (*34)* |
| IN77-20-1-1 | *S. italica* | India | 1977 | (*34)* |
| KANSV1-4-1 | *S. viridis* | Japan | 1975 | (*34)* |
| NI913 | *S. viridis* | Japan | 1974 | (*34)* |
| US71 | *Setaria* sp. | USA | ND | (*56*) |
| SZEC1-1-1 | *Eleusine coracana* | Japan | 1978 | (*34)* |
| GFEC1-5-1 | *El. coracana* | Japan;Gifu | 1977 | (*34)* |
| UG77-7-1-1 | *El. indica* | Uganda;Serere | 1977 | (*34)* |
| UG77-17-1-1 | *El. coracana* | Uganda, Kabanyolo | 1977 | This study |
| UG77-15-1-1 | *El. coracana* | Uganda;Kabanyolo | 1977 | (*34)* |
| CD156 | *El. indica* | Ivory Coast | 1989 | (*56*) |

| | | | | |
|---|---|---|---|---|
| Z2-1 | *El. coracana* | Japan | 1977 | (*57*) |
| MZ5-1-6 | *El. coracana* | Japan;Miyazaki | 1976 | (*30*) |
| NP10-17-4-1-3 | *El. coracana* | Nepal | 1975 | *(34)* |
| NI1006 | *El. africana* | Japan | 1975 | *(34)* |
| NI1011 | *El. boranensis* | Japan | 1975 | *(34)* |
| IN77-36-1-1 | *El. indica* | India | 1977 | *(34)* |
| NI986 | *Eragrostis lehmanniana* | Japan, Kumamoto | 1975 | *(34)* |
| FSECu1-1-1 | *Er. curvula* | Japan, Fukushima | 1988 | *(34)* |
| SZECu1-1-1 | *Er. curvula* | Japan, Shizuoka | 1988 | *(34)* |
| Br35 | *Urochloa plantaginea* | Brazil | 1990 | *(30)* |
| TP1 | *Lolium perenne* | Japan;Tochigi | 1997 | *(34)* |
| TP2 | *L. perenne* | Japan;Tochigi | 1997 | *(30)* |
| TP3 | *L. perenne* | Japan;Tochigi | 1997 | This study |
| AK1 | *L. perenne* | Japan;Akita | 1998 | *(34)* |
| AK2 | *L. perenne* | Akita, Japan | 1998 | This study |
| LW1 | *L. perenne* | Japan | 1999 | This study |
| LW3 | *L. perenne* | Japan;Yamanashi | 1999 | *(34)* |
| FI5 | *L. perenne* | Japan;Chiba | 1998 | *(34)* |
| LpKY97 | *L. perenne* | U.S.A. | 1997 | *(56)* |
| Br58 | *Avena sativa* | Brazil | 1990 | *(30)* |
| Br2 | *Triticum aestivum* | Brazil | 1990 | *(34)* |
| Br3 | *T. aestivum* | Brazil | 1990 | *(34)* |
| Br5 | *T. aestivum* | Brazil | 1990 | *(34)* |
| Br8 | *T. aestivum* | Brazil | 1990 | *(34)* |
| Br46 | *T. aestivum* | Brazil | 1990 | *(34)* |
| Br48 | *T. aestivum* | Brazil | 1990 | This study |
| Br49 | *T. aestivum* | Brazil | 1990 | *(34)* |
| Br50 | *T. aestivum* | Brazil | 1990 | *(34)* |
| Br108.1 | *T. aestivum* | Brazil | 1992 | *(34)* |
| Br115.7 | *T. aestivum* | Brazil | 1992 | *(34)* |
| Br115.12 | *T. aestivum* | Brazil | 1992 | *(34)* |
| Br116.5 | *T. aestivum* | Brazil | 1992 | *(30)* |
| Br118.2 | *T. aestivum* | Brazil | 1992 | *(30)* |
| Br126.1 | *T. aestivum* | Brazil | 1992 | *(34)* |

| | | | | |
|---|---|---|---|---|
| Br127.1 | *T. aestivum* | Brazil | 1992 | (*34)* |
| Br127.11 | *T. aestivum* | Brazil | 1992 | (*34)* |
| Br130.8 | *T. aestivum* | Brazil | 1992 | (*34)* |
| Br130.9 | *T. aestivum* | Brazil | 1992 | (*34)* |
| BTMP-2(b) | *T. aestivum* | Bangladesh | 2017 | This study |
| BTMB-5(b) | *T. aestivum* | Bangladesh | 2017 | This study |
| BTGP-6(e) | *T. aestivum* | Bangladesh | 2017 | This study |
| B71 | *T. aestivum* | Bolivia | 2012 | (*58)* |
| ZMV18_06 | *T. aestivum* | Zambia | 2018 | Open Wheat Blast* |
| ZMV19_09 | *T. aestivum* | Zambia | 2019 | Open Wheat Blast* |
| ZMV20_03 | *T. aestivum* | Zambia | 2020 | Open Wheat Blast* |

*http://openwheatblast.net/

### Table S4. Primers used in this study

| Objectives | Name | F (Forward primer, 5' -> 3') | Description |
|---|---|---|---|
| Mapping of *Rmo2* | AGT11-F | GTGGGAGGGTGGGTGTTAG | CAPS marker |
| | AGT11-R | CTGGAGATCCTCCCCAAAC | |
| | AGT18-F | GCTTGCAGCTGACTTCCTCT | Presence/absence marker (presence in R81) |
| | AGT18-R | TGGAGCGAAGACATCACTTG | |
| | AGT62-F | CCACCAGTAGGGCTCAGAAG | CAPS marker (ApaI) |
| | AGT62-R | CTCCATGAGCTTTCCTCAGC | |
| | AGT107-F | GAGACCCGCCATACCATCAG | CAPS marker (BstUI) |
| | AGT107-R | AGGTTATCAATGCGTGGGCA | |
| | AGT113-F | GTGGCCACCACGACCAATTA | Presence/absence marker (presence in R81) |
| | AGT113-R | GCATGCCCTCTAGATGCTGAT | |
| | AGT116-F | ATGAGGGAAATGCCATGGGG | CAPS marker (PsiI) |
| | AGT116-R | TCCTCGTGCTCGTACTCTGA | |
| | AGT119-F | CACGACATCTCCCACACCAT | CAPS marker (XhoI) |
| | AGT119-R | CACGGATCATCCTCACCCAG | |
| | AGT122-F | TCAGCATTGCGAACATTGCC | CAPS marker (PsiI) |
| | AGT122-R | TACCGGCCATCGCAACATTA | |
| | AGT125-F | CCTTGAAGATGATGCGGTGC | CAPS marker (DpnII) |
| | AGT125-R | GTCACCTTCGTCAACACCGA | |
| | ABC1037-2-F | CCCTATTCGTGCATGCCTCT | CAPS marker (TaqI) |
| | ABC1037-2-R | GGCTGCTACAGGGATTTGGT | |
| | AGT143-F | TCGAATGGGTGGACATGCAA | CAPS marker (Hpy99I) |
| | AGT143-R | TGGCCCGAATTGTGGTACTC | |
| | DN4-F | CTTTGGTGGGAGGAAACGGA | CAPS marker (Hpy99I) |
| | DN4-R | GTTTCCATCACCACGTTGCC | |
| Cloning of *Rmo2* | InF-pBUH3_1037d-F | GCTCACTAGTGGATCATGGCGGCTGCCAGTAAG | for In-Fusion cloning of *Rmo2.d* |
| | InF-pBUH3_1037d-R | GCTTCCCGGGGGATCTCAAGTGAGGAAGGTCTTC | |
| | InF-pBUH3_1037a-F | GCTCACTAGTGGATCATGGCGGCTGCCAGGAAG | for In-Fusion cloning of *Rmo2.a* |
| | InF-pBUH3_1037a-R | GCTTCCCGGGGGATCATATGGCAATTGCTGCAACT | |
| Mapping of *Rwt7* | Ta7D_7-F | ACCAGGTGGACGGAACTCAACTC | CAPS marker (HhaI) |
| | Ta7D_7-R | CCAGTACTCCTCTACGCCATTGTTC | |

| | | | |
|---|---|---|---|
| | Ta7D_18-F | AACCTGGGGAAATGGACCAATGAG | CAPS marker (Hpy166) |
| | Ta7D_18-R | CAGCAACAGTGAGTTCTCTGCATAAG | |
| | Ta7D_23-F | CACGGCGTGTCCGGGGCATATT | CAPS marker (SacI) |
| | Ta7D_23-R | ACGCCCTCGATGTCGTACCAGTGCT | |
| | Ta7D_31-F | TATTTATCTTCGTGCTGGCGGTCAC | CAPS marker (HhaI) |
| | Ta7D_31-R | AGTTGAATTTTAAGTTCCCCCGACAG | |
| | Ta7D_38-F | CTACCTTGACCGCACACTCTTATGTG | CAPS marker (NotI) |
| | Ta7D_38-R | GGTGCGCGACGTAGGCTAGATTG | |
| | Ta7D_61-F | GTTTCACCGGCCTAAACTAGTCCTTC | CAPS marker (PsiI) |
| | Ta7D_61-R | CTGGCTTTGTATTACGCTTCCTAGCC | |
| | Ta7D_87-F | AATGGCGGCCTTTAACTGACGTATAG | CAPS marker (AfaI) |
| | Ta7D_87-R | TATGCGCTCCCCTGAGAAACTATTTG | |
| | CS7D-Rmo2h-F | AGCCATTTGCTTGCTTGACAGC | CAPS marker (SspI) |
| | CS7D-Rmo2h-R | CGACAGGCAATCAGTCCC | |
| | barc184-F | TTCGGTGATATCTTTTCCCCTTGA | SSR marker |
| | barc184-R | CCGAGTTGACTGTGTGGGCTTGCTG | |
| | cdf41-F | TAAAGTCTCAGGCGACCCAC | SSR marker |
| | cdf41-R | AGTGATAGACGGATGGCACC | |
| | gdm145-F | TGAAGGACAAATCCCTGCAT | SSR marker |
| | gdm145-R | TCCCACCTTTTTGCTGTAGA | |
| | cfd31-F | GCACCAACCTTGATAGGGAA | SSR marker |
| | cfd31-R | GTGCCTGATGATTTTACCCG | |
| Cloning of *Rwt7* | InF-pBUH3_Rwt7-F | GCTCACTAGTGGATCATGAAGCAAAAGATTGTGGTG | for In-Fusion cloning of *Rwt7* |
| | InF-pBUH3_Rwt7-R | GCTTCCCGGGGGATCTTACGTTCTCCTTGTAATGC | |
| Mapping of *PBY2* | MGM43-F | TGCATGAAGCTGATTTGCTC | SSR marker |
| | MGM43-R | TTTGACTCTCGCTCCCTCTC | |
| | MGM188-F | TGGGAAGTCGATAGTCAGGAA | SSR marker |
| | MGM188-R | TGCACGATTAGCTGGTGAAG | |
| | MOSSR2_1-F | GGTCTGTTGGAGGGTTTTCG | SSR marker |
| | MOSSR2_1-R | ATCACCGAGCTTGACATTGC | |
| | POTS50_2-F | TTGATGTGCAAGCTCAGGTC | Presence/absence marker (presence in Br48) |
| | POTS50_2-R | GATGGCAAGCAGTGAAGTGA | |
| | POTS65_1-F | GAGCATACCGGCCTTTGTC | Presence/absence marker (presence in Br48) |
| | POTS65_1-R | CCCCAAATCATATTCGTCCA | |

| | | | |
|---|---|---|---|
| | POTS57-F | CTTCTTCACTCGCCCAAAAG | Presence/absence marker (presence in Br48) |
| | POTS57-R | GGAAATAAACGCTCCCATGA | |
| | MGM47-F | GGCAGGTAGACACGACCAAG | SSR marker |
| | MGM47-R | CTTCCGGATGAATCACCAAC | |
| | MGM48-F | GCCGCAACAATGACTTAACA | SSR marker |
| | MGM48-R | CCGCGGTAGGTAAATGAGAG | |
| | MGM51-F | GTAACCAGGCCGTTTCAAGA | SSR marker |
| | MGM51-R | GGAGGTTGCAGAAGGACAGA | |
| | POTC94118-F | GCCTCATCGAGACTTTCAGG | Presence/absence marker (presence in Br48) |
| | POTC94118-R | CCATTCGTGGTAAACCCATC | |
| | POTC98605-F | AAGGCATTTGTCGCTGTACC | Presence/absence marker (presence in Br48) |
| | POTC98605-R | ATTTGGCAATTTTGGAAACG | |
| | POTC40320-F | CCGGTGTGGGTAAATGAGAC | Presence/absence marker (presence in Br48) |
| | POTC40320-R | CGGGAATGCCTTTATCTTGA | |
| | POTC98875-F | GGTGTGGTGATGGAAATGAA | Presence/absence marker (presence in Br48) |
| | POTC98875-R | GCGAAAGCGGGAGAAAAC | |
| | POTC98650-F | AAACCCCGTCGTTATTTTCC | Presence/absence marker (presence in Br48) |
| | POTC98650-R | CAAGGTCTTTTGGACCCGTA | |
| | POTC97724-F | GGGAATCTCTTATCGCCACA | Presence/absence marker (presence in Br48) |
| | POTC97724-R | TAGTTCGCCAATGCGTGTTA | |
| Cloning of *PBY2* | ORF40-cl-F | CTTGCAGTGTCCGCAAGAAC | for cloning of *PBY2* and *PBY1* |
| | ORF40-cl-R | CCTGAAGAGCGGACACCTTT | |
| Disruption of *PBY2* | disPBY2_F1-F | ATCGATAAGCTTGATTCCTTGCACCTATTTCTACT | for amplification of downstream of *PBY2* |
| | disPBY2_F1-R | TAGTGTCACCTAAATTATTTGGCCAAAAAAGTTCAAG | |
| | disPBY2_F2-F | ATTTAGGTGACACTATAGAAC | for amplification of HPH gene |
| | disPBY2_F2-R | TTAACGACAGGTTTTTAATACGACTCACTATAGGG | |
| | disPBY2_F3-F | AAAACCTGTCGTTAAACCAG | for amplification of upstream of *PBY2* |
| | disPBY2_F3-R | CTGCAGGAATTCGATATTTCTCGGCGTGACAGC | |
| | HY | GGATGCCTCCGCTCGAAGTA | for split marker method |
| | YG | CGTTGCAAGACCTGCCTGAA | |
| Cloning of *Rpg1* and *WTK4* | Rpg1_Mrx_F | TTGCCTTCCACGTACTTTCC | for cloning of *Rpg1* |
| | Rpg1_Mrx_R | CTTCTGGGGTGTCACCACTT | for cloning of *Rpg1* |
| | Wtk4-CDS-F2 | CCCTGGTGTTGGTTTGATTT | |

| | | | |
|---|---|---|---|
| | Wtk4-CDS-R3 | GTCCTTCTTCGCTGTGCTG | |
| Protoplast assay | InF-dSP-PWT7-F | TGTGTGTGCAGATCGATGCGAAGATGCGTGGTTG | for In-Fusion cloning of *PWT7* without its signal peptide |
| | InF-dSP-PWT7-R | GGAAATTCGAGCTCGTTAATTTTCCCACCGTGTTC | |
| | InF-dSP-PBY2-F | TGTGTGTGCAGATCGATGAAGAAACCCGAGGAATG | for In-Fusion cloning of *PBY2* without its signal peptide |
| | InF-dSP-PBY2-R | GGAAATTCGAGCTCGGTCATTGTTCAACCTGCGTG | |
| | InF-H2-F | TGTGTGTGCAGATCGATGGCGGCTGCCAGTAAG | for In-Fusion cloning of *Rmo2.d, Rwt7*, and *Rmo2.d*::HMA(*Rwt7*) |
| | InF-H2-R | CTCCTGGCCGCTGTCGTACTTTGACGCGGGCCC | |
| | InF-H7-F | TGTGTGTGCAGATCGATGAAGCAAAAGATTGTGGTG | |
| | InF-H7-R | CTCGAACGTAAGTTCGTCTTTCTTCTTGATGATCTC | |
| | InF-P2-F | GAACTTACGTTCGAGTTCTT | |
| | InF-P2-R | GGAAATTCGAGCTCGTCAAGTGAGGAAGGTCTTC | |
| | InF-P7-F | GACAGCGGCCAGGAGGAG | |
| | InF-P7-R | GGAAATTCGAGCTCGTTACGTTCTCCTTGTAATGC | |
| | InF-Rpg1-Mrx-F | TGTGTGTGCAGATCGATGATGCGAAGGATGGTGATG | for In-Fusion cloning of *Rpg1* |
| | InF-Rpg1-Mrx-R | GGAAATTCGAGCTCGTCATTTTACGTTCCTCTTCAACTTC | |
| | InF-Wtk4-ORF-F1 | TGTGTGTGCAGATCGATGAAGAAGATCGTGCTGAA | for In-Fusion cloning of *WTK4* |
| | InF-Wtk4-ORF-R1 | GGAAATTCGAGCTCGTCATGGCGTCTGCTGCAG | |
| Swapping HMA domain & barley transformation | InF-H7P2-HindIII-F | ATCCCCCGGGAAGCTATGAAGCAAAAGATTGTGGTG | |
| | InF-H7P2-HindIII-R | AGCTGGTACCAAGCTTCAAGTGAGGAAGGTCTTC | |
| Sequencing *Rmo2* alleles | ABC1037-seq-F1 | TGTTGACTTGATGGCGGCTG | for cloning of *Rmo2* alleles & sanger sequencing |
| | ABC1037-5-F1 | ATTTTGCCTTCCACTTTCCT | for cloning of *Rmo2* alleles & sanger sequencing |
| | ABC1037-5-F2 | AGATCTATTTTGCCTTCCAC | for cloning of *Rmo2* alleles & sanger sequencing |
| | ABC1037-seq-R1 | CAGCCAATGCTTCAAGTGAGGA | for cloning of *Rmo2* alleles & sanger sequencing |
| | ABC1037-seq-F2 | GCTGGATAAGGACATGATGC | for sanger sequencing |
| | ABC1037-seq-R2 | CCTCATTCGCCAGTTTTCAC | for sanger sequencing |
| | ABC1037-seq-R3 | TAGAGCTATTCGGATGCATG | for sanger sequencing |
| | ABC1037_590_F | GGAGGAAGCCTTGA | for sanger sequencing |
| | 1037-PK1-seq-out-R1 | GGCATCATGTCCTTATCC | for sanger sequencing |
| | ABC1037_1160_seq_F | CCAACTATGTTCTGCTCAGAAA | for sanger sequencing |
| Distribution analysis of *Rmo2* and *Rpg1* | $PK^{Rpg1F\text{-}1}$ | ATAATCTTATGAGGGTCCGA | |
| | $PK^{Rpg1R\text{-}1}$ | CTCGATTGGACTCCACGCA | |
| | $HMA^{Rpg1F\text{-}2}$ | TTGCCTTCCACGTACTTTCC | |
| | $PK^{Rpg1R\text{-}2}$ | CTTCTGGGGTGTCACCACTT | |
| | $HMA^{Rmo2F\text{-}3}$ | TGTTGACTTGATGGCGGCTG | |

| | | | |
|---|---|---|---|
| | $PK^{Rpg1R-3}$ | CTCGATTGGACTCCACGCA | |
| Expression analysis | HvActin-F | TCGCTCCACCTGAGAGGAAG | internal control in barley |
| | HvActin-R | GCTAGGATGGACCCTCCGAT | |
| | Hv-GAPDH_cw1 | CGTTCATCACCACCGACTAC | internal control in barley |
| | Hv-GAPDH_ccw1 | CAGCCTTGTCCTTGTCAGTG | |
| | Exp-Rmo2-4F | AATTGTGCAGACACCCCTCC | for *Rmo2* expression analysis |
| | Exp-Rmo2-4R | ATTGTAGGCCTTCGGGTTCG | |
| | Rpg1_Ex3_cw2 | GCCGGTGTACTATCCCTTTC | for *Rpg1* expression analysis |
| | Rpg1_Ex4_ccw2 | TGTCGGACCCTCATAAGATT | |
| | CDCP-L | CAAATACGCCATCAGGGAGAACATC | internal control in wheat |
| | CDCP-R | CGCTGCCGAAACCACGAGAC | |
| | Exp-Rwt7-3F | GTTGACAGCAGTCGCCGTTC | for *Rwt7* expression analysis |
| | Exp-Rwt7-3R | ACAACCTTCGCAGCTTTCCC | |
